# Beyond defense: Glucosinolate structural diversity shapes recruitment of a metabolic network of leaf-associated bacteria

**DOI:** 10.1101/2023.12.04.567830

**Authors:** Kerstin Unger, Ali K. Raza, Teresa Mayer, Michael Reichelt, Johannes Stuttmann, Annika Hielscher, Ute Wittstock, Jonathan Gershenzon, Matthew T. Agler

## Abstract

Leaf bacteria are critical for plant health, but little is known about how plant traits control their recruitment. Aliphatic glucosinolates (GLSs) are secondary metabolites present in leaves of Brassicaceae plants in genotypically-defined mixtures. Upon damage, they are broken down to products that deter herbivory and inhibit pathogens. Using two *A. thaliana* genotypes with different aliphatic GLS profiles, we find that structural variants differentially affect commensal leaf bacteria: In the model genotype Col-0, GLS breakdown products (mostly from 4-methylsulfinylbutyl-glucosinolate) are potentially highly toxic to bacteria but have no effect on natural leaf colonization. In contrast, in an *A. thaliana* genotype from a wild population, GLS (mostly allyl-GLS) enriches Burkholderiales bacteria, an effect also detected in nature. Indeed, *in-vitro* as a carbon source, intact allyl-GLS specifically enriches a Burkholderiales-containing community in which Burkholderiales depend on other bacteria but in turn increase community growth rates. Metabolism of different GLSs is linked to breakdown product detoxification, helping explain GLS structural control of community recruitment.

## Introduction

Plant health depends to a great extent on microbial colonization of roots and leaves (*1*). Besides pathogens which are detrimental to plant health, other microbes also play important roles in plant fitness. Non-pathogenic bacteria, especially, are important for protecting plants against microorganisms that can cause disease. For example, non-pathogenic bacteria enable plant survival upon germination in soil in the presence of potentially detrimental soil fungi (*2*) and can protect leaves against pathogen attack (*3*, *4*). Thus, it is important to understand which factors determine the colonization of bacteria in organs like leaves. In this context, both plant-microbe and microbe-microbe interactions are relevant. While we have a good understanding of 1-on-1 interactions between pathogens and plants in leaves, it is mostly unknown how non-pathogenic bacteria survive there, and in turn which host traits shape their assembly into communities in naturally colonized plants.

In order to successfully colonize leaves, all bacteria need to overcome several hurdles, such that the taxa that finally reach the surface and endosphere of plant leaves have been filtered by several factors (*5*, *6*). First, to find their way onto or into the leaf, microbes must overcome physical hurdles such as low water availability (*7*) and regulated stomatal openings (*8*). Next, they need to evade the plant immune system (*9*, *10*), which is made up of sensors of microbial molecular patterns (pattern-triggered immunity and effector-triggered immunity) as well as an arsenal of defensive secondary metabolites. Finally, the leaf environment is thought to be oligotrophic and very heterogeneous (*11*, *12*), making it a challenge to find nutrient sources.

The plant immune system, especially, is thought to play important roles in selection and regulation of bacterial colonizers. For example, flagellin proteins of bacteria are finely tuned to evade pattern-triggered immunity (*13*), and generation of oxidative stress is important for regulating opportunistic pathogens (*14*). On the other hand, little is known about how the diversity of secondary metabolites, which can contribute to defense, shape leaf colonization. To study this, we focused on the well-known glucosinolate-myrosinase system and asked how it might influence leaf bacterial communities of healthy *Arabidopsis thaliana* plants. Glucosinolates (GLSs) are secondary metabolites produced by plants in the Brassicaceae and related families. They share a common backbone structure consisting of a β-D-glucopyranose residue linked via a sulfur atom to a (Z)-N-hydroximinosulfate ester with variable side chains (*15*). The chemical diversity of GLSs is determined by their side chains which result from their biosynthesis from different amino acids (*16*). Aliphatic GLSs are a diverse group of GLSs derived from methionine, alanine, leucine, isoleucine or valine, whereas indole or benzenic GLSs are synthesized from aromatic amino acids (*15*). In *A. thaliana*, the plant genotype defines the ability to synthesize a certain set of aliphatic GLSs, but the precise GLS mixtures are controlled developmentally, organ-specifically and in response to environmental factors. Wild genotypes isolated across Europe are typically characterized by a single major leaf aliphatic GLS that defines a “chemotype” (*17*).

Aliphatic GLSs are constitutively present particularly in epidermal cells and in specialized cells along the vascular bundles (*18*). In addition, up to 5% of total leaf aliphatic GLSs may be present on the leaf surface (*19*). Although considered biologically inactive, GLSs can be activated upon leaf damage by myrosinases, which hydrolyze the glucose moiety leading to rearrangement to various breakdown products, including isothiocyanates (ITCs), nitriles, and epithionitriles. The final chemical mixture depends on the aliphatic GLS structure and the presence of plant specifier proteins (*20*), as well as on abiotic conditions like temperature and pH (*21*).

The role of aliphatic GLSs and their breakdown products have been best studied with respect to their defensive role against herbivorous insects. However, ITCs especially are well-known for antimicrobial properties against a broad range of plant and human pathogens *in vitro* (*22*, *23*) and in the model *A. thaliana* genotype Col-0, they help protect against bacterial and fungal pathogens (*24*, *25*). In turn, microbial pathogens have adapted to the Brassicaceae with mechanisms to deal with toxic breakdown products such as detoxification and efflux pumps encoded by *sax* (<u>s</u>urvival in *<u>A</u>rabidopsis* e<u>x</u>tracts) genes to cope with ITC stress during infection (*24*, *25*). While ITCs have long been considered to be present only after activation upon plant cell damage, there is now evidence for a constant turnover of GLSs to ITCs and cysteine as part of sulfur-cycling in plants (*26*). Indeed, 4MSOB-ITC was detected in the apoplastic fluids of healthy Col-0 leaves (*27*) and the low concentrations present were reported to be enough to affect *P. syringae* virulence. Similarly, GLSs and their breakdown products exuded from roots are known to affect rhizosphere bacteria community assembly (*28*) and fumigation of soils with ITCs or bulk biomass from Brassicaceae plants suppresses detrimental microorganisms in soils (*29*, *30*). Therefore, we hypothesize that aliphatic GLSs and their breakdown products might function as a filtering mechanism for bacterial leaf colonization of healthy plants. Given the wide chemical differences among aliphatic GLSs and their breakdown products in different plant genotypes, we further reasoned that different GLSs may shape the leaf bacterial community in distinct ways.

## Results

### Wild *A. thaliana* populations from Jena have distinct aliphatic GLS profiles

We studied five distinct, wild populations of *A. thaliana* located in Jena, Germany (Fig. 1A, 1C, Tab. S1). We had previously isolated individual plants from these populations, grown them in the laboratory, and characterized their leaf GLS profiles (*31*). The chemotype of all the isolates differed from that of the model genotype Col-0, where 4MSOB-GLS is the principal aliphatic GLS (Fig. 1B). The main GLS in three out of five isolates was 3-hydroxypropyl GLS (3OHP-GLS) (SW1, JT1, PB). In one isolate (NG2) allyl-GLS dominated, and another isolate (Woe) produced both 2-hydroxy-3-butenyl GLS (2OH3But-GLS) and allyl-GLS. In 2022 and 2023 we additionally analyzed the GLS profiles of NG2 and Woe plants sampled directly from wild populations. We found that Woe GLSs were the same as those previously extracted from this population, but wild NG2 contained both 2OH3But-GLS and allyl-GLS, similar to Woe (Fig. S1). Since plants of all these Jena populations possessed a completely different aliphatic GLS composition than the widely used reference genotype Col-0, we compared one of them, NG2, to Col-0 to understand how GLS diversity affects the assembly of leaf bacterial communities.

**Figure 1.**
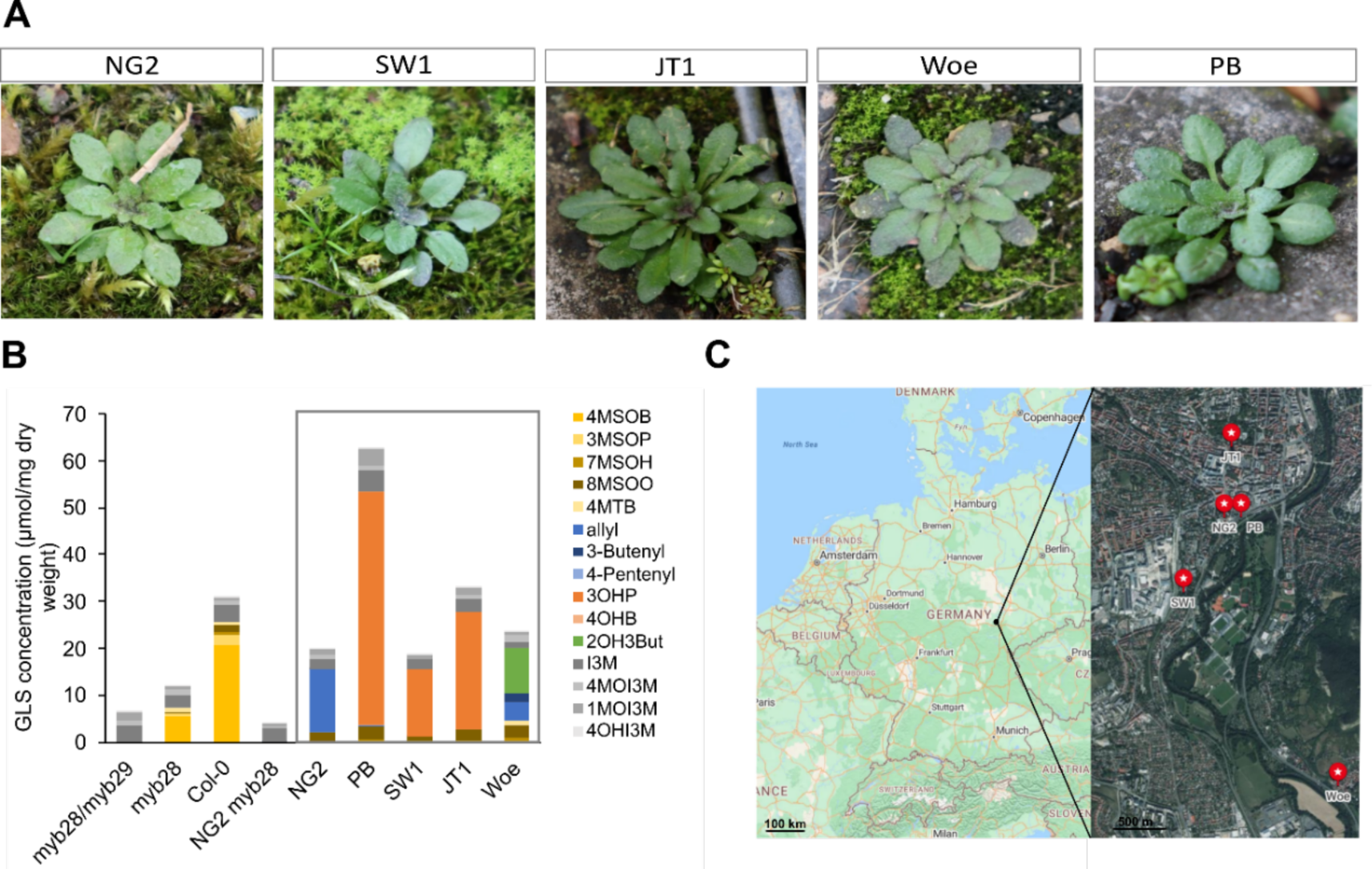
Local *A. thaliana* populations in Jena produce distinct GLS profiles. We used these differences to study the impact of these leaf metabolites on bacterial community composition. (**A**) Individual *A. thaliana* plants of the five selected populations in Jena, Germany: NG2 (Neugasse), SW (Sandweg), JT1 (Johannistor), Woe (Wöllnitz), PB (Paradiesbahnhof) in February 2022. (**B**) Average GLS concentrations of 3-4 replicates of the five local *A. thaliana* populations (grey box), the reference genotype Col-0 and respective aliphatic GLS mutants in Col-0 and NG2 background. Colored GLSs are aliphatic, gray shades are indole GLSs. Abbreviations for GLSs are listed in the methods. (**C**) Map of Germany and Jena showing the sampling locations of the five populations (created with Microsoft Bing Maps), additional information is available in Tab. S1.

### Aliphatic GLS breakdown products of certain *A. thaliana* genotypes inhibit growth of commensal leaf bacteria

We assumed that inhibition of bacterial growth would be the most likely mechanism by which aliphatic GLSs or their breakdown products would shape bacterial leaf communities. To compare toxicity between GLS-derived products in *A. thaliana* Col-0 and NG2, we homogenized leaves to mix the GLSs with myrosinases and release GLS breakdown products into the medium. We prepared what we refer to as “leaf extract medium” from the isolated genotype NG2 (which mainly produces allyl-GLS) and the reference genotype Col-0 (mainly 4MSOB-GLS), and from two transgenic lines, Col-0 *myb28* (with reduced aliphatic GLSs, Fig. 1B) and Col-0 *myb28/myb29* (with no aliphatic GLSs, Fig. 1B). We tested the leaf extract media against 100 diverse bacterial isolates recovered from *A. thaliana* leaves collected from the wild populations NG2 and PB (Data S1, Fig. S2). For most isolates, growth in Col-0 leaf extract medium was poor. Isolates of *Curtobacterium* spp., *Xanthomonas* spp. and *Pseudomonas* spp. all grew slightly better with aliphatic GLS-reduced *myb28* leaf medium, and all tested genera grew better with aliphatic GLS-free *myb28/myb29* leaf medium indicating growth inhibition by aliphatic GLS breakdown products. Interestingly, NG2 leaf extract medium was less inhibitory, and several strains grew significantly more than in Col-0 *myb28/myb29* extract (Fig. 2A). These opposing effects suggest that aliphatic GLS breakdown products from different *A. thaliana* genotypes act very differently towards bacteria.

**Figure 2:**
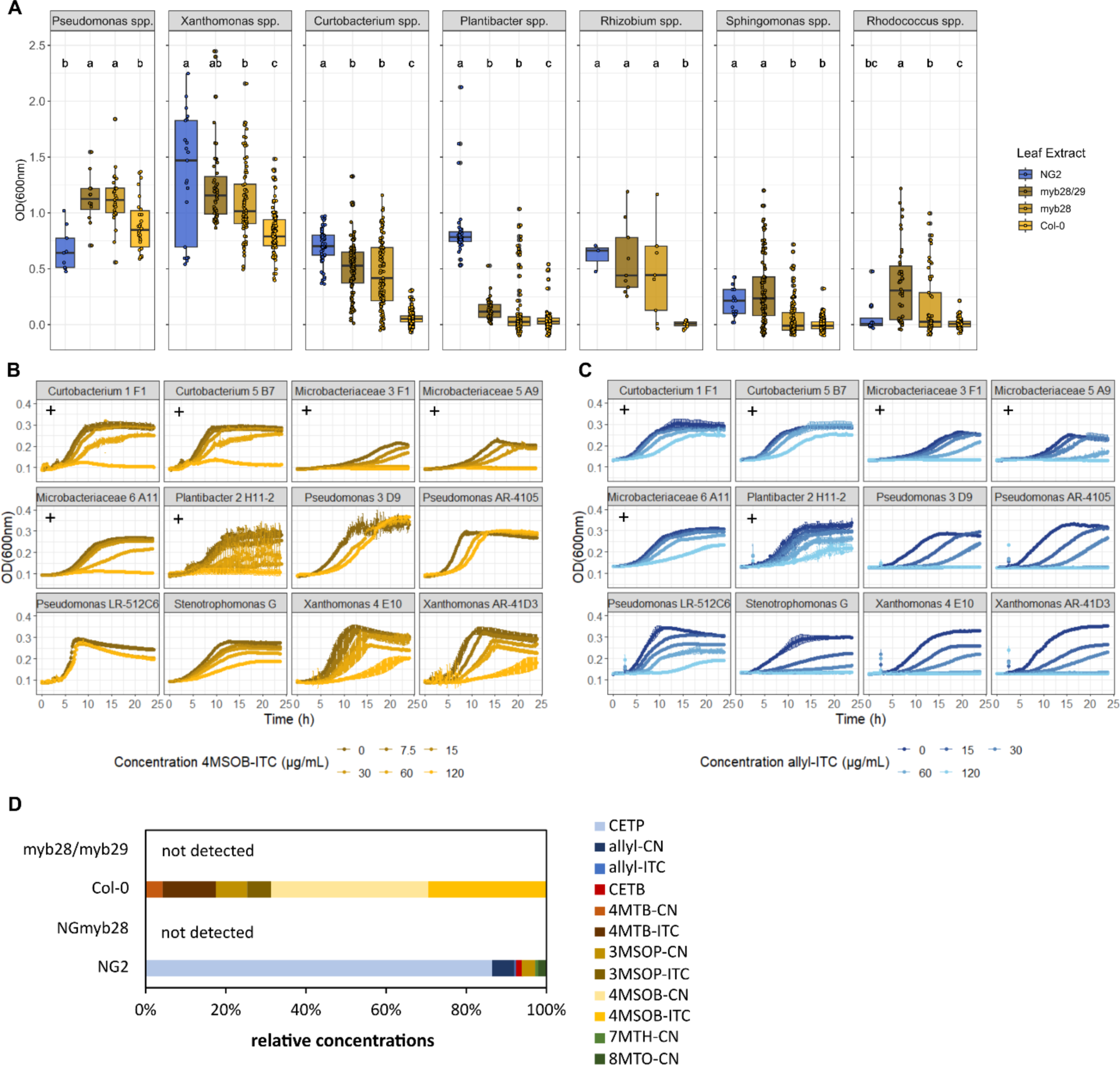
Effects of GLS degradation products on growth of diverse leaf colonizing bacteria. (**A**) Final OD_600_ of bacterial strains grown in leaf media of different *A. thaliana* genotypes and mutants. Each strain was measured in three technical replicates and at least in two leaf extract media. Some strains were tested repeatedly, and data of several strains was agglomerated at genus level for better visibility. Number of strains per genus: 7 *Pseudomonas*, 12 *Xanthomonas*, 16 *Curtobacterium*, 13 *Plantibacter*, 2 *Rhizobium*, 37 *Sphingomonas*, 13 *Rhodococcus* spp. (individual plots: Fig. S2) Letters indicate statistical significance based on ANOVA followed by a Tukey post hoc-test with alpha = 0.05. (**B,C**) Growth curves of a set of 12 bacterial strains in R2A medium supplemented with 4MSOB-ITC (**B**) or allyl-ITC (**C**). Gram-positive strains are marked with a +, the remaining strains are gram-negative. Mean and standard deviation of three replicates are shown per condition. (**D**) Relative concentration of aliphatic GLS breakdown products in NG2, NG*myb28*, Col-0 and *myb28/myb29* leaf homogenates per gram fresh weight. The average of three replicates per genotype is shown. CN = nitrile, ITC = isothiocyanate; additional details on the abbreviations for GLS breakdown products are listed in the methods section.

Assuming that ITCs would be the main inhibitory compounds in leaf extract medium, we tested pure ITCs against the bacterial isolates. As expected, 4MSOB-ITC inhibited growth of most isolates, especially gram-positive strains, consistent with the broad inhibitory effect of Col-0 leaf extract. Gammaproteobacteria like *Stenotrophomonas* sp., *Xanthomonas* and *Pseudomonas* spp. were more resistant (Fig. 2B). For the strains of the latter two genera this greater resistance corresponds to the presence of *sax* genes which are known ITC resistance genes and for *Pseudomonas syringae* strains to the ability to degrade 4MSOB-ITC (Fig. S3, Tab. S2). Allyl-ITC was also toxic, especially to gram-negative colonizers in apparent contradiction to the results with the NG2 leaf extract where these bacteria grew well (Fig. 2C). Upon investigation of the actual GLS breakdown products, we found that Col-0 leaf homogenates contained high levels of 4MSOB-ITC (29.6 ± 6.1%) and 4MSOB-CN (39.3 ±7.4%), the corresponding nitrile. In NG2 homogenates, however, the predominant aliphatic GLS breakdown product was not the corresponding ITC, but rather the epithionitrile 3,4-epithiobutanenitrile (CETP, 86.6 ± 2.1%), known to be derived from allyl-GLS by the action of an additional plant specifier proteins (*20*). Only a minor proportion of allyl-GLS was converted to the corresponding nitrile (allyl-CN, 5.4 ± 0.7%) and even less to allyl-ITC (0.3 ± 0.0%) (Fig. 2D). Thus, this epithionitrile, although found in leaf homogenates at high levels, apparently hardly inhibited the growth of most bacterial isolates. Together, both aliphatic GLS structure and the types of breakdown products influence toxicity towards leaf colonizing bacteria.

### Aliphatic GLSs do not decrease bacterial colonization, but rather increase colonization of specific taxa in the NG2 genotype

Based on the apparent higher toxicity of Col-0 leaf homogenates, we reasoned that there would be higher potential for GLSs to affect leaf bacterial community assembly in Col-0 than in NG2 plants. To test both genotypes together, we knocked out *myb28* in the NG2 background, which completely eliminated aliphatic GLSs in the leaves of this genotype (Fig. 1B). Then, we grew both genotypes and their respective aliphatic GLS-free mutants in natural soil collected in Jena and performed 16S rRNA gene amplicon sequencing to characterize the bacterial community in surface-sterilized (mostly endophytic bacteria) and whole (including all surface bacteria) leaves. Importantly, we used hamPCR (*32*) so that results are normalized to single-copy host gene abundance and reflect differences in absolute bacterial abundances, not just relative abundances.

We did not find any significant differences in alpha or beta diversity of endophytic or total leaf bacterial communities between Col-0 and its aliphatic GLS-free mutant *myb28/myb29* (Data S2, Fig. S4), agreeing with previous work (*27*). In NG2, we also did not observe differences in endophytic communities (Data S2, Fig. S4), but the beta diversity of total leaf communities was significantly affected by the *myb28* knockout (Fig. 3B, PERMANOVA: R2=0.14704 p=0.042 Jaccard; R2=0.2472 p=0.065 Bray-Curtis). To our surprise, NG2 also had *higher* total bacterial loads in leaves compared to NG*myb28* (Wilcoxon test, p=0.032, Fig. 3A). Further, a differential abundance analysis of taxa showed that 13 of 14 genera that were affected by genotype were *enriched* in NG2 compared to NG*myb28* (Fig. 3C). Seven of these 13 belonged to the order Burkholderiales: two from the family *Methylophilaceae* and five from *Oxalobacteriaceae*. The other taxa enriched in NG2 WT leaves were diverse but included members of the Rhizobiales and the Flavobacteriales. Together, the results refuted our hypothesis that the toxicity of GLS breakdown products affected commensal leaf colonization of healthy plants. Instead, the enrichment of bacteria in NG2 with aliphatic GLSs compared to NG*myb28* suggests a strong positive effect of allyl-GLS, but not 4MSOB-GLS on specific taxa.

**Figure 3:**
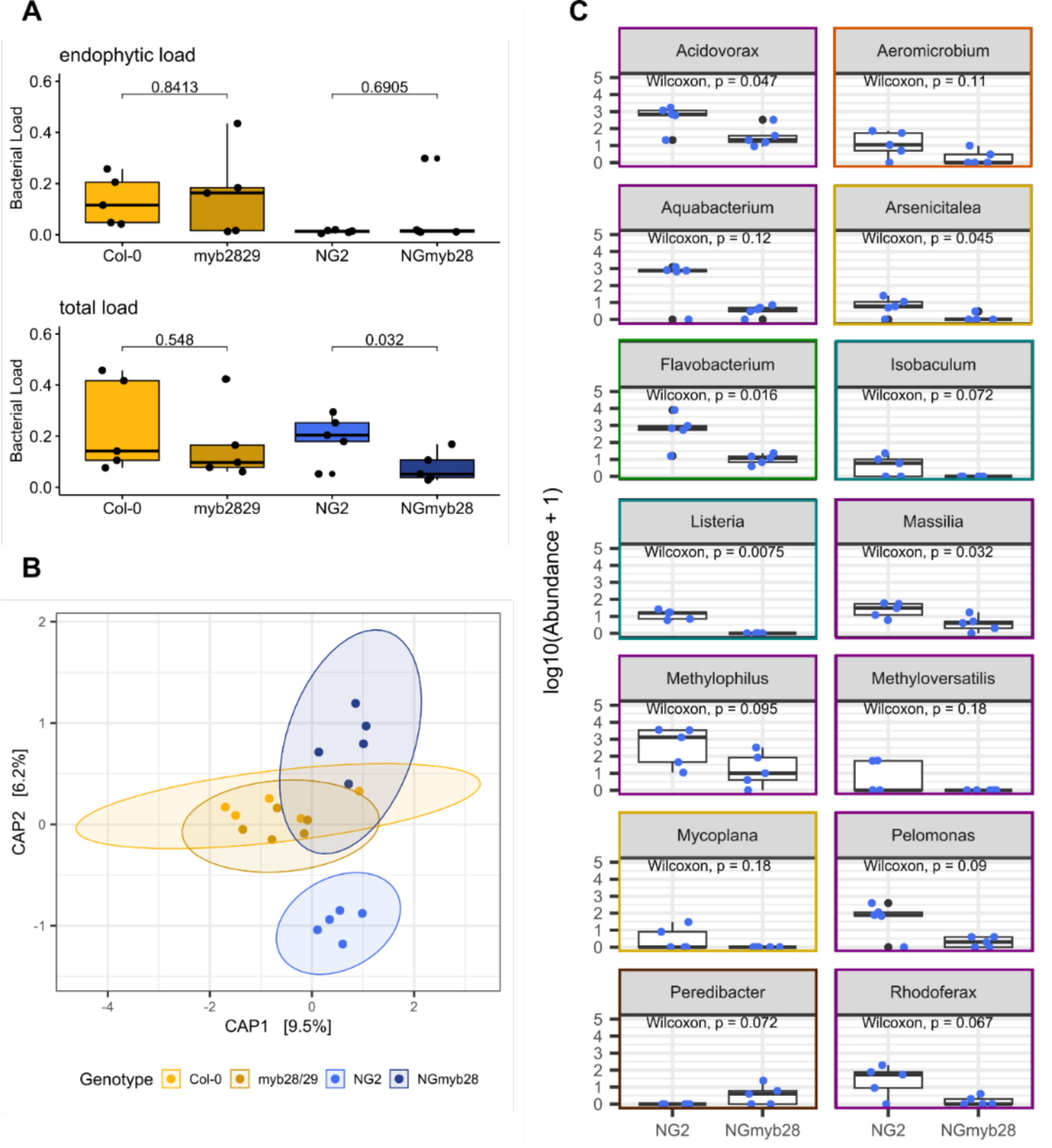
Bacterial community analysis of leaves with and without aliphatic GLSs in NG2 and Col-0 background. Bacterial community composition of leaves of 3-week-old plants was assessed by amplicon sequencing of 16S rRNA genes (n=5). (**A**) Bacterial loads of total and endophytic leaf communities of *A. thaliana* assessed by normalization to plant GI reads, pairwise Wilcoxon test was used to check for significances. (**B**) Beta diversity of total leaf communities visualized as constrained PCoA of Jaccard index. Pairwise comparisons (PERMANOVA between NG2-NG*myb28*, Col-0-*myb28/myb29*) revealed significant differences between NG2-NG*myb28*. (**C**) Differentally abundant taxa on NG2 compared to NG*myb28* leaves using DESeq analysis with a cutoff of alpha = 0.05. Significant taxa were log10-transformed and plotted with pairwise Wilcoxon tests on the abundances. Colors of the boxes show the family level: purple = Burkholeriales, brown = Bacteriovoracales, yellow = Rhizobiales, blue = Lactobacilliales, green = Flavobacteriales, orange = Propionibacteriales.

### A. thaliana *leaves in the wild specifically enrich taxa associated with aliphatic GLSs*

To test whether the bacterial taxa enriched in the lab *in planta* in response to aliphatic GLSs are also enriched in *A. thaliana* in the wild, we sampled leaves of *A. thaliana* together with leaves of other sympatric, ground-dwelling ruderal plants growing in our five wild populations (Fig. 1) and characterized the whole leaf bacterial communities. Bacterial communities associated with *A. thaliana* leaf samples were overall similar to those of other plants. However, 2.8% of the variation in community composition corresponded significantly to the plant type (*A. thaliana* vs. other plants), and plant type also interacted with other variables, especially location (6.6% of variation) (Fig. 4A). Differential abundance analysis revealed that the *A. thaliana*-specific signature involved several taxa, some of which were enriched in *A. thaliana* leaves both at the NG2 location alone and across all locations (Fig 4B, Fig. S5). Among these, several Burkholderiales genera were prominent, including *Acidovorax*, *Rhizobacter*, *Rhodoferax*, *Variovorax* and the methylotroph *Methylotenera*, as well as *Flavobacterium* (Flavobacteriales). This list parallels the differential enrichment seen in NG2 vs. NG*myb28* in the lab colonization experiments, which included *Flavobacterium*, *Acidovorax*, *Rhodoferax* and other apparently methylotrophic Burkholderiales (Fig 3C). Apart from *Methylotenera,* no taxa associated with aliphatic GLSs or NG2, most notably Burkholderiales, were enriched in *A. thaliana* samples taken from the Woe population, even though these plants share a very similar chemotype with NG2 (Fig. S1, Fig. S5). On the other hand, populations PB and SW1, both with a 3OHP-GLS chemotype, were still enriched in similar Burkholderiales (Fig. 1B, Fig. S5). Thus, while taxa associated with aliphatic GLSs were generally enriched in most of these wild populations of *A. thaliana* studied, other genotypic or ecological factors shape their recruitment locally.

**Figure 4:**
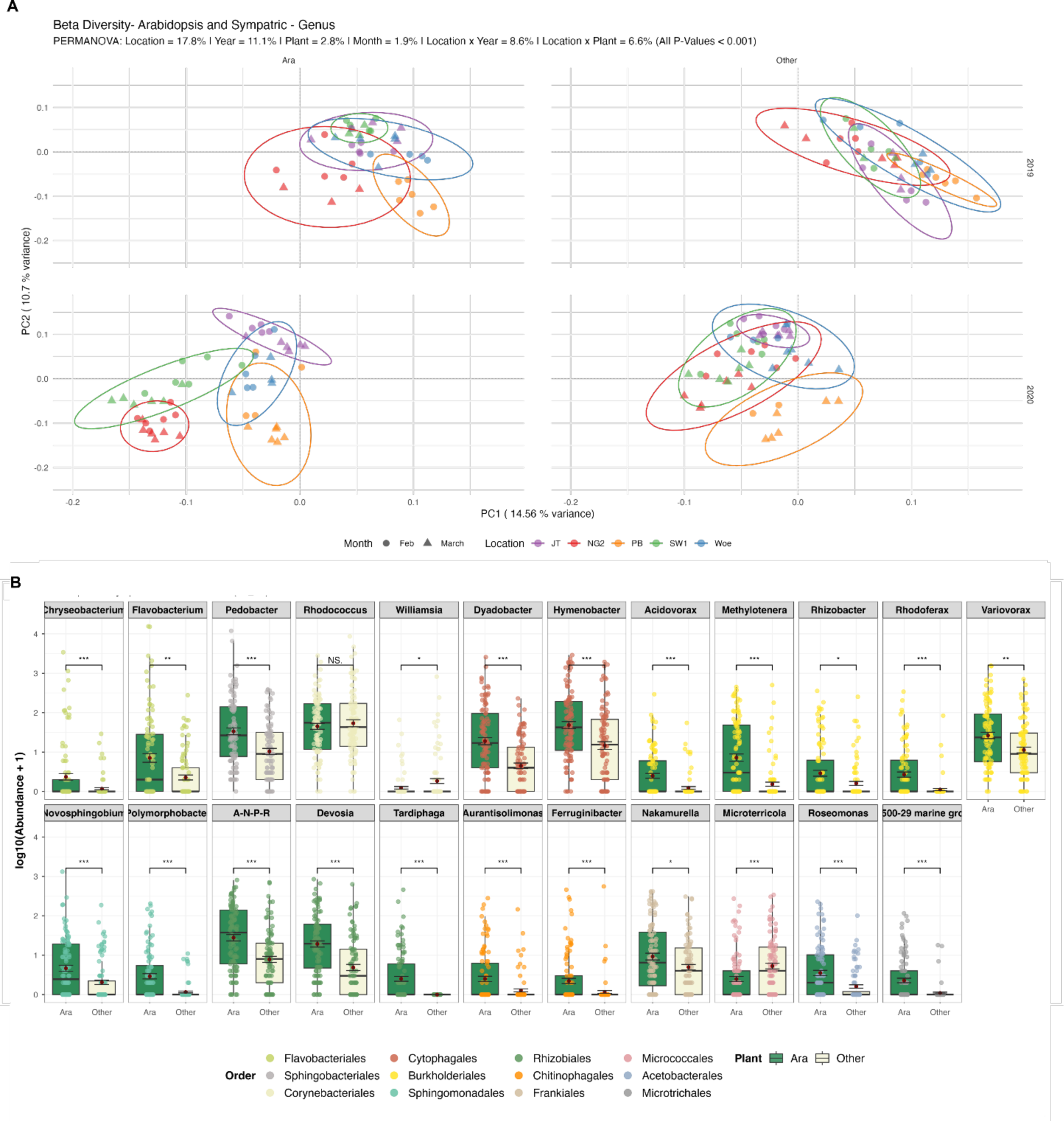
Leaf bacterial community signature of *A. thaliana* compared to sympatric plants across years and locations. Leaf bacterial community compositions in five locations (NG2, JT, PB, Woe, SW1) in February and March of 2019 and 2020 assessed by amplicon sequencing of 16S rRNA genes. **(A)** PCoA of leaf bacterial communities based on Aitchison distances from centered log ratio (CLR)-transformed genus-level compositions. Points represent individual samples, categorized by location (color) and month (shape). 95% confidence intervals are shown for each location-year combination via ellipses. PCoA was performed on all data together and the plots are facetted by year and plant type. Variance explained is indicated on the axes. The fraction of variance explained by the factors are shown based on PERMANOVA (P < 0.001). (**B**) Differentially abundant taxa using DESeq analysis with a cutoff of alpha = 0.05. Significant taxa were log10-transformed and plotted with p-values calculated using the Benjamini-Hochberg method. Dark red points indicate the mean abundance. The overlaid jitter points represent individual samples and are colored by order.

### With allyl-GLS as the sole carbon source, bacterial communities are enriched in Burkholderiales

We hypothesized that bacteria on the leaf surface of the NG2 *A. thaliana* population may utilize surface allyl-GLS as a carbon source, leading to the enrichment of certain strains and overall higher bacterial loads on NG2 plants compared to NG*myb28*. To test this, we washed bacteria from leaves of wild NG2 plants to inoculate M9 minimal medium supplemented with allyl-GLS, 4MSOB-GLS or glucose as the sole carbon source. Nitrogen and sulfur, also found in GLSs, were not limited in the base medium. An additional experimental trial supplemented with glucose followed by allyl-GLS (Fig. 5A). The leaf surface wash contained 5.43 x 10^6^ CFU/mL. We enriched for three passages, where each time 10% of the volume was transferred to a new substrate so that any remaining leaf carbon sources would have been insignificant (∼1000x diluted).

**Figure 5:**
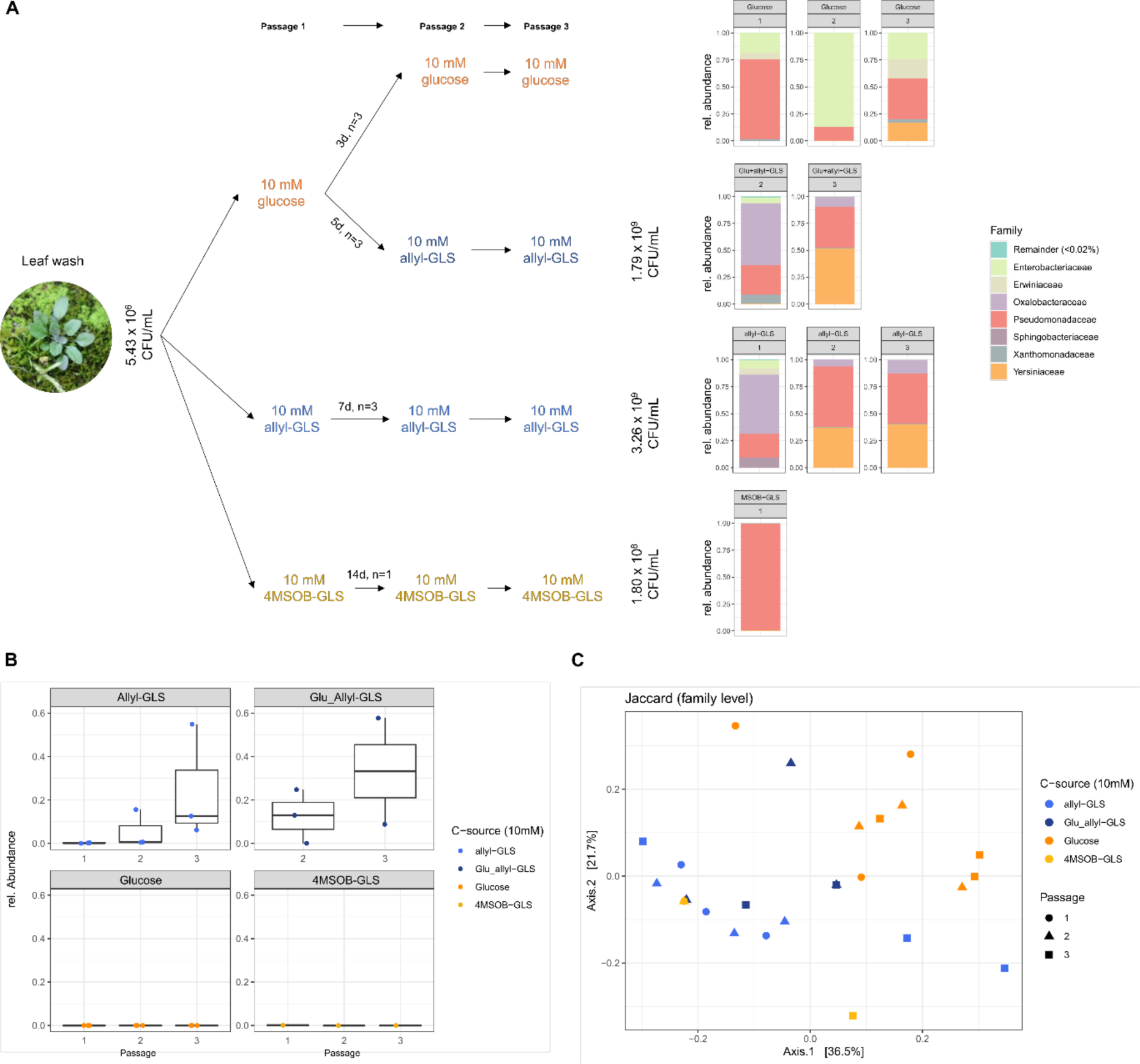
Enrichment of bacterial strains from NG2 leaf surface on different aliphatic GLSs as sole carbon source. (**A**) Schema of enrichment process with initial and final CFUs, growth intervals and number of technical replicates. Bar charts show the community composition after the third passage assessed by 16S rRNA gene amplicon sequencing. The charts show data agglomerated on family level. Families below 0.02% relative abundance were merged and classified as “Remainder”. Only replicates with >100 reads were considered. (**B**) Relative abundance of *Oxalobacteraceae* family over the three passages in all enrichments. (**C**) Beta diversity measured by Jaccard distances of all enrichments on family level with significant differences based on C-source (PERMANOVA: p=0.006, R2=0.257).

Passage intervals were adjusted for growth rate differences. It took seven days for growth on allyl-GLS, 14 days on 4MSOB-GLS, three days on glucose, and five days on glucose followed by allyl-GLS. By the final passage, bacterial populations reached an average of 3.26 x 10^9^ CFU/mL on allyl-GLS-supplemented medium, 1.80 x 10^8^ CFU/mL on 4MSOB-GLS, and 1.79 x 10^9^ CFU/mL on glucose followed by allyl-GLS (Fig. 5A).

16S rRNA gene amplicon sequencing showed that the final communities grown on glucose- or 4MSOB-GLS-supplemented medium were dominated by *Pseudomonadaceae*. On 4MSOB-GLS *Pseudomonadaceae* made up almost 100% of the total, whereas both *Enterobacteriaceae* and *Pseudomonadaceae* were abundant on glucose. The communities growing on allyl-GLS were distinct and showed one of two different configurations, each with at least one member of the order Enterobacterales, one Burkholderiales (always a *Janthinobacterium* ASV, belonging to *Oxalobacteraceae*) and one *Pseudomonadaceae*, regardless of whether there was a pre-enrichment on glucose. In three of five replicates, one *Yersiniaceae* ASV (Enterobacterales) dominated together with *Pseudomonadaceae* and *Oxalobacteraceae* (Fig. 5A). In the other two replicates, a community developed that was dominated by *Oxalobacteraceae* and *Pseudomonadaceae* but also included *Enterobacteriaceae* and *Erwiniaceae* (both Enterobacterales). ASVs belonging to *Oxalobacteraceae* increased in relative abundance over the course of the passages, suggesting that they increased in importance in the communities over time and that community formation was dynamic (Fig. 5B). In general, the carbon source was correlated to 25.7 % of the variation between communities (p=0.006, PERMANOVA, Jaccard) (Fig. 5C). In conclusion, when the carbon source was allyl-GLS, the community was enriched in Burkholderiales, a group which was also associated with allyl-GLS *in planta* both in lab-grown plants and in wild populations.

### Only a *Yersiniaceae* strain metabolized aliphatic GLSs, with rates depending on the structure

We next isolated bacteria from the communities growing on medium with 4MSOB-GLS and allyl-GLS as sole carbon sources to determine which individual taxa can directly utilize GLSs. The recovered taxa closely reflected the taxa identified by amplicon sequencing (Tab. S3, Fig. 5A). From the 4MSOB-GLS medium, four *Pseudomonas* isolates were recovered, but none grew successfully on 4MSOB-GLS within six days (Fig. S6A). This makes sense given the 14-d growth time required to reach a relatively low cell density in the passages on 4MSOB-GLS medium, and underscores that bacteria grow only very slowly on 4MSOB-GLS. From the allyl-GLS medium, seven isolates representing most of the taxa detected at family level were tested, but only one *Yersiniaceae* strain (top BLAST hit: *Rahnella,* hereafter R3) grew on allyl-GLS within six days (Fig. S6B, Fig. 6A). R3 metabolized an average of 69.1 % of the allyl-GLS in the medium in six days of growth. Metabolite analysis of the culture supernatant showed that very little of the allyl-GLS was recovered as allyl-ITC (0.006 ± 0.001 mM), but that on average 93.1% was metabolized to the presumably less toxic breakdown product allyl-amine (11.6 ± 4.7 mM) (Fig. 6B). We also tested whether R3 could grow on 2OH3But-GLS, which wild NG2 plants produce in leaves together with allyl-GLS. It did, but the lag-phase was longer (Fig. 6A) and only 20.9 % of 2OH3But-GLS was consumed within nine days. In contrast to allyl-GLS, little of the 2OH3But-GLS was converted to the corresponding amine (0.03 ± 0.01 mM), but instead ∼58.9 % was found as goitrin (1.77 ± 0.33 mM) which results from spontaneous cyclization of the unstable 2OH3But-ITC (Fig. 6B).

**Figure 6:**
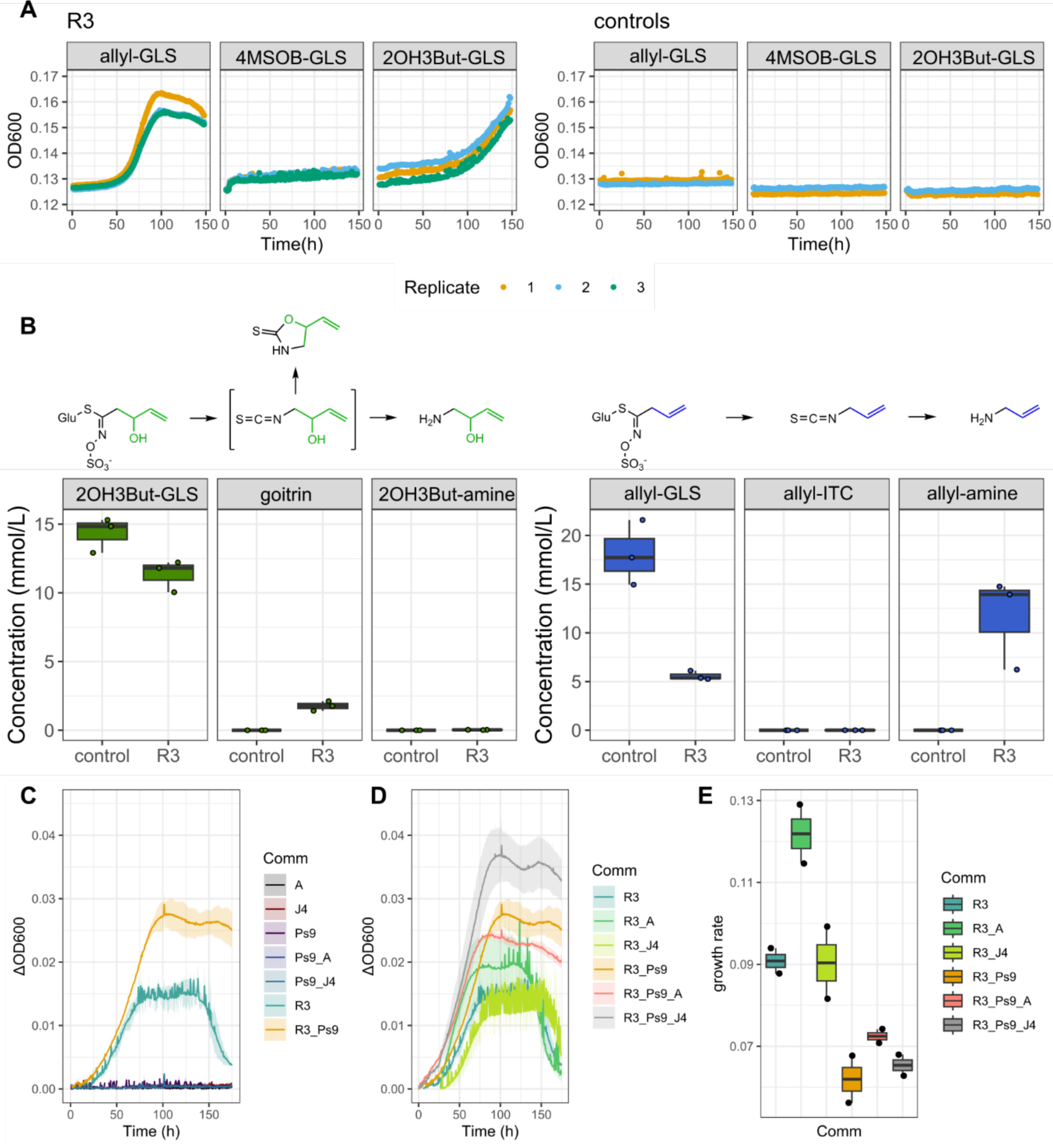
Growth on and utilization of diverse aliphatic GLSs by bacterial strains recovered from the enrichments. (**A**) R3 growth in M9 medium supplemented with 10 mM allyl-GLS, 4MSOB-GLS or 2OH3But-GLS. OD_600_ was measured every hour, water served as negative control (n=3). (**B**) Analysis of 2OH3But-GLS, allyl-GLS and degradation products in R3 inoculated medium after six days (allyl-GLS) and nine days (2OH3But-GLS) of incubation (n=3). **(C,D**) Growth curves of mixtures of bacterial strains in M9 medium supplemented with 10 mM allyl-GLS, after pre-culturing the same communities twice in the same medium for seven and five days. OD_600_ was measured every hour, the first OD measurement was subtracted of all following ones to blank (n=2). (**E**) Growth rates calculated based on the curves in C,D.

### Interactions with Burkholderiales within communities develop to shape growth dynamics on allyl-GLS

The previous results suggested that if allyl-GLS was primarily used as a carbon source in leaves, Burkholderiales would not be directly enriched. We reasoned, however, that enrichment could be indirect if the growth of R3 on allyl-GLS as a carbon source could support the growth of other taxa. Therefore, we combined R3 with other strains to observe how growth would be affected. When co-cultivated in a single six-day passage, R3 with *Pseudomonas* Ps6 or Ps9 reached a higher maximum OD_600_ than R3 alone, suggesting more efficient carbon utilization. Combining R3 with other taxa had no effect or reduced total growth (Fig. S6C, S6D).

Next, we designed an experiment in which all possible combinations of R3, *Janthinobacterium* J4 and *Pseudomonas* Ps9 (representatives of the taxa that were always enriched on allyl-GLS, Fig. 4B) were passaged three times on allyl-GLS. We also included combinations in which J4 was replaced by *Acidovorax* 4E11-1 (hereafter A4), a representative of a Burkholderiales taxa enriched on both lab and wild plants containing aliphatic GLSs. As expected, by the third passage, we only observed growth in communities where R3 was present (Fig. 6C), confirming that this strain has unique roles in mobilizing resources from GLSs when GLS is the sole carbon source. As previously observed, the addition of Ps9 to R3 resulted in higher maximal OD_600_ compared to R3 monocultures (Fig. 6C, 6D). A4 together with R3 resulted in a far higher growth rate than any other strain combination and a slightly increased OD_600_, whereas the addition of J4 to R3 gave similar growth compared to the R3 monoculture (Fig. 6D). Among the tripartite cultures, adding J4 to the R3/Ps9 mix resulted in earlier growth compared to R3/Ps9 alone as well as the highest observed OD_600_ (Fig. 6D, 6E). R3/Ps9/A4 resulted in similar increases in growth rate but a slightly decreased OD_600_ compared to R3/Ps9 alone (Fig. 6D, 6E). Together, the experiments demonstrate that the growth of R3 on allyl-GLS can support diverse taxa and these taxa in turn shape the fitness of the community. In particular, Burkholderiales tend to positively influence community growth, helping to explain why these bacteria are consistently associated with allyl-GLS in leaves, despite not metabolizing it themselves.

## Discussion

Plant exudates are well-described to shape assembly of root and rhizosphere microbial communities (*33*, *34*) by serving as nutrient sources for rhizosphere bacteria (*35*), inhibiting growth of certain taxa to protect the plant (*36*, *37*) or altering microbial physiology and activity (*38*) and microbial interactions (*39*). In leaves, however, few compounds are definitively known to positively recruit specific bacteria: Lab experiments have suggested that sugars non-specifically recruit leaf bacteria (*40*), while in the wild, positive recruitment of methylotrophic bacteria is known based on simple carbon compounds like methanol (*41*, *42*) that are thought to be by-products of plant metabolic processes (*11*). Our findings show that recruitment in leaves may be even more prevalent and that secondary plant metabolites such as aliphatic GLSs can be important in orchestrating leaf colonization of commensal bacteria (Fig. 7).

**Figure 7:**
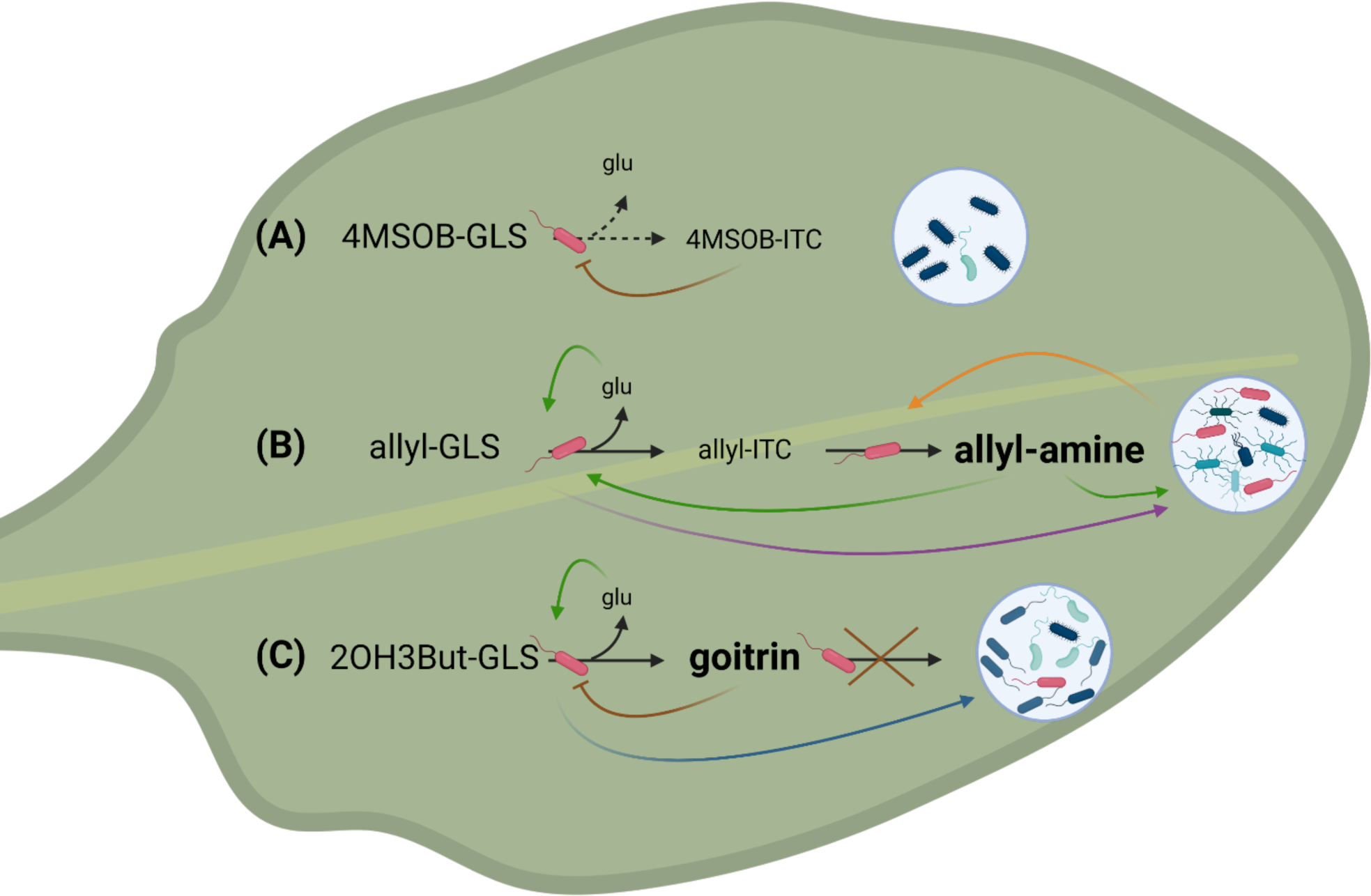
Hypothesis for mechanisms of community assembly dependent on GLS utilization and metabolic feedback loops. The font size depicts the concentrations of GLS breakdown products, where the main products are visualized in bold. We propose three scenarios which would result in different bacterial communities: (**A**) 4MSOB-GLS is rarely utilized because 4MSOB-ITC inhibits bacteria that can hydrolyze this GLS. (**B**) Allyl-GLS hydrolysis to allyl-ITC, which can be relatively easily detoxified to allyl-amine by the hydrolyzing bacteria. Hydrolysis of the GLS makes glucose available, and this and/or cross-feeding possibly involving allyl-amine (green arrow) and other secreted metabolic byproducts (purple arrow) leads to a positive feedback loop promoting more rapid GLS degradation and growth of the community. In return, metabolic products from the community can promote growth of the GLS utilizer (yellow arrow). (**C**) 2OH3But-GLS can be hydrolyzed to some degree, and thus the resulting glucose moiety promotes some growth. However, the resulting breakdown product goitrin likely either has inhibitory effects or is not accessible, reducing metabolic activity of the hydrolyzing bacteria and the potential for metabolic byproducts (blue arrow) to surrounding bacteria. Schematic drawing was created with BioRender.com.

Given the well-known antimicrobial effects of aliphatic GLS breakdown products (*22–25*), it may seem surprising that this defence system did not negatively impact the colonization of *A. thaliana* leaves by commensal bacteria. However, the lack of a suppressive effect of aliphatic GLSs on non-pathogenic bacteria in healthy Col-0 leaves is supported by previous work (*27*). In our work, aliphatic GLSs were even found to promote the recruitment of specific bacterial taxa. The leaf surface is a challenging environment for microbes because nutrients and water are limited. While no single nutrient clearly limits bacterial growth in leaves more than others, carbon is especially well-studied and known to be highly patchily distributed (*12*). However, plant secondary metabolites are present (*11*) and these could potentially be used to fill nutritional deficiencies experienced by leaf bacteria. Aliphatic GLSs are known to be present on the leaf surface (*19*), and other non-defensive roles are known for example in butterfly oviposition behaviour, whereby adult cabbage butterflies (*Pieris rapae*) express gustatory receptors in their tarsi that sense allyl-GLS to identify host leaves for oviposition (*43*). From a microbial perspective, every GLS molecule contains a glucose moiety, which can be enzymatically cleaved by myrosinases to yield glucose. In addition, this hydrolysis reaction releases sulfate (S), and the further metabolism of breakdown products could provide more carbon or nitrogen in the form of amines. Supporting nutritive roles, human gut bacteria break down GLSs (*44*), and myrosinase-producing bacteria have been identified in both soil and plant roots (*45*, *46*) as well as in the phyllosphere (*47*). GLSs as resources are costly, however, since myrosinase cleavage also releases an aglycone that can rearrange to form ITCs or other toxic products. Thus, resistance to ITCs and other GLS breakdown products is also a requirement for efficient utilization of GLSs.

Here, we investigated allyl-GLS primarily as a carbon source and isolated only one strain, R3, that was able to survive on minimal medium with aliphatic GLSs as the only carbon source. R3 degraded the GLSs, but its ability to do so was strongly linked to the GLS structure, which can help explain why different *A. thaliana* GLS chemotypes caused strikingly different effects on leaf bacterial community formation. The aliphatic GLSs of Col-0 had no effect on leaf bacterial community composition and *in vitro* growth on 4MSOB-GLS was slow and relatively poor. This could be because it was not degraded by the bacterial myrosinase, which are known to show substrate specificity (*45*, *48*). The success of R3 might also depend on formation and degradation of ITCs during bacterial GLS metabolism. ITCs are directly toxic to bacteria (*49*) affecting activity and growth (*27*), which potentially results in a negative feedback loop (Fig. 7). However, ITC hydrolases can convert ITCs to non-toxic amines (*24*), as seen with R3, and they also exhibit substrate specificities (*50*), which can explain why this strain grew best on allyl-GLS (Fig. 7B). R3 also grew on 2OH3But-GLS, but more slowly, likely because goitrin was the main product (Fig. 7C). Goitrin results from spontaneous ring formation of the unstable 2OH3But-ITC (*51*), making the ITC group unavailable for further detoxification (*52*).

In natural *A. thaliana* populations and *in planta* in the lab we did not observe high abundances or consistent enrichment of Yersiniaceae in contrast to *in vitro* enrichments. In addition, although wild *A. thaliana* produced mixes including 2OH3But-GLS (NG2) or mainly 3OHP-GLS (PB/SW) that are likely to be more difficult to access than allyl-GLS, they still apparently enriched Burkholderiales taxa that were associated with allyl-GLS in the lab. Assuming this enrichment is indeed due to GLS metabolism, it is possible that bacteria that can access GLSs do not need to be present in high abundances so that we did not detect Yersiniaceae. On the other hand, many bacteria can utilize GLSs (often other Enterobacterales) (*46*) and myrosinases have been suggested to be enriched in the phyllosphere (*47*). Given the differing specificities of myrosinases and ITC hydrolases for substrate structures, it is very likely that other bacteria may be able to functionally replace Yersiniaceae *in planta*, preventing its consistent enrichment in the phyllosphere.

Although we only definitively identified one bacterial strain, R3, that could grow on allyl-GLS, it was enriched together with a substantially more diverse community. Burkholderiales bacteria especially were consistently enriched with R3 and associated with allyl-GLS in both lab and wild plants. Burkholderiales are important for plants, being linked to growth promotion (*53*) and antifungal properties (*54*). In *A. thaliana*, they contribute to suppression of pathogenic fungi, which is required for survival when seedlings germinate in soil (*2*). Therefore, understanding their enrichment may lead to ways to promote plant health via microbiomes. We hypothesize that metabolic cross-feeding from R3 likely contributed to the growth of other taxa (Fig. 7). Cross-feeding can support diverse communities on small numbers of primary metabolites (*55*) and we previously found that leaf bacteria are exceptionally prepared and adaptable to cross-feeding (*56*). GLS metabolism in particular would result in an abundance of breakdown products like glucose or allyl-amine that may be valuable carbon or nitrogen resources for co-occurring taxa.

In bacterial communities associated with R3 growing on aliphatic GLSs, metabolic interactions appear to have a reciprocal nature, especially in the way Burkholderiales taxa had the capacity to increase efficiency of community growth. While the mechanism of this improvement is unclear, Burkholderiales are metabolically complex and have often been observed as part of consortia degrading complex compounds (*57*). In enrichments of bacteria from the surface of peppers grown on capsaicin as the sole carbon and nitrogen source, only a combination of a Burkholderiales (*Variovorax*) with a *Pseudomonas* strain could grow, probably by reciprocal exchange of capsaicin-derived carbon and nitrogen (*58*). Additionally, some Burkholderiales have the capacity to fix nitrogen in the phyllosphere (*59*, *60*), and this could contribute to overall bacterial growth on surface GLSs.

A supportive role of Burkholderiales in the growth of other commensal leaf bacteria is consistent with previous work. In different wild *A. thaliana* populations, we identified a *Comamonadaceae* (Burkholderiales) genus as a “hub”, highly positively correlated to the abundance of other bacteria in leaves of wild *A. thaliana* (*61*). Later work in the same population using abundance-weighted networks similarly found a tightly positively correlated module of *Comamonadaceae* and other bacteria, suggesting these bacteria increase and decrease in abundance together (*62*). Thus, we hypothesize that Burkholderiales play key roles in leaf metabolic networks, at least in part including GLS-based carbon economies, and future work should be aimed at dissecting these roles.

Aliphatic GLSs and their breakdown products are some of the best-studied leaf secondary metabolites involved in defence against pathogens. For example, 4MSOB-ITC protects against non-host pathogens by direct antimicrobial effects (*25*) and by suppressing expression of the type III secretion system, a major virulence factor of the pathogen *Pseudomonas syringae* (*27*). Diverse other ITCs also are known to inhibit plant and human pathogens (*22*, *23*). This has likely led to selective pressure resulting in pathogen specialization for GLS-containing plants: *Pseudomonas syringae* DC3000, *Pectobacterium* spp. and the fungal pathogen *Sclerotinia sclerotiorum* all express *sax* genes which enable virulence in the presence of ITCs in *A. thaliana* and cabbage plants (*24*, *25*, *63*). Accordingly, we found *sax* gene homologs in genomes of the opportunistic pathogens *Pseudomonas* and *Xanthomonas,* even though they were isolated from healthy leaves. *Pseudomonas* 3D9 and AR-4105 possess the ITC hydrolase SaxA and were able to degrade 4MSOB-ITC. On the other hand, most leaf-colonizing bacterial taxa were strongly inhibited by GLS breakdown products like 4MSOB-ITC in Col-0. Thus, even though ITCs did not shape colonization of healthy leaves as we and others observed (*27*), they would be released in large amounts during herbivory or due to necrotic pathogens, probably strongly re-shaping leaf bacterial communities. That might explain why herbivory increased *P. syringae* bacterial loads in leaves of *Cardamine cordifolia* (*64*), since *P. syringae* are likely to have *sax*-gene mediated tolerance to ITCs. Additionally, defence responses including the release of ITCs might also occur in apparently healthy tissues due to small-scale responses (*59*) to local attack by opportunistic pathogens found in these leaves (*65*). On the other hand, events leading to GLS breakdown would likely have less effect in NG2 leaves, where allyl-GLS breaks down to a presumably less toxic epithionitrile. Therefore, understanding how interactions between leaf-damaging organisms and microbiomes together affect plant fitness (*64*) will require both focusing in on localized, small-scale effects (*66*) and looking beyond the model *A. thaliana* genotype Col-0, into diverse other chemotypes.

There are several important directions for future work to enable possible applications of these findings. Especially, it is necessary to evaluate how particular GLSs and/or GLS mixtures shape recruitment in nature. GLSs clearly play dual roles in recruitment and defence and our results suggest these roles will probably vary depending on leaf bacteria in a community context. Thus, to fully understand GLS roles in nature, it will be necessary to evaluate how effects in controlled lab-conditions are shaped by factors like both the biotic and abiotic environment. We were also so far unable to measure allyl-GLS consumption directly in leaves because there are still significant uncertainties about how to feed leaf bacteria *in-planta* with specific metabolites in a controlled but realistic way. For example, it is unclear how surface GLS become available to bacteria and how realistic leaf localization can be artificially reproduced. However, metabolite localization together with feeding and tracing experiments would contribute to a better understanding of GLS-mediated community assembly by revealing GLS turnover rates and helping elucidate how breakdown products shape activity of the leaf bacterial community. At any rate, the finding that assembly of communities on aliphatic GLSs is plant chemotype-specific and thus defined by plant genomes sets the stage for developing new approaches to shape and maintain balance in plant leaf microbiomes.

## Material and Methods

### Local *A. thaliana* populations in Jena

We worked with the widely used reference *A. thaliana* Col-0, its single and double knock-out mutants *myb28* and *myb28/myb29* (*67*) and the local genotype NG2. The latter was identified and isolated in spring 2018 as one out of five wild *A. thaliana* populations in Jena, Germany: NG2, PB, SW1, JT1, and Woe (*31*). One individual plant of each population was propagated in the lab from a single seed for two generations to generate relatively uniform, homozygous lines for further experiments. The isolated plants are available under the names Je-X from NASC (Tab. S1).

### Knock-out of *myb28* in local NG2 A. thaliana

To study effects of aliphatic GLSs in NG2 we generated an aliphatic GLS-free mutant in NG2 background by knocking out the Myb28 transcription factor using a genome editing procedure by an RNA-guided SpCas9 nuclease (*68*). The plasmid pDGE347 was programmed for six target sites within MYB28 (AT5G61420; AAAAAACGTTTGATGGAACA<u>GGG</u>; TTCAAATTCTCATCGACCGT<u>AGG</u>; GATCGGGAGTATTGCTTGT<u>CGG</u>; GCTTCTAGTTCCAACCCTA<u>CGG</u>; GAAACCATGTTGCAACTGGA<u>TGG</u>; GAAACGTTTCTTGCAACTCA<u>AGG</u>). The respective plasmid (pDGE816) was transformed into *Agrobacterium tumefaciens* strain GV3101 pMP90 and plants of accession NG2 were transformed by floral dipping as previously described (*69*). Floral dipping resulted in CRISPR-guided transformation events already in the germ cells of the plant and therefore T1 generation seeds were screened for successfully transformed seeds (indicated by RFP expression in seeds) (*70*). Primary transformants and non-transgenic individuals from the T_2_ population were PCR screened and Sanger sequenced to isolate homozygous *myb28* lines using oligonucleotides myb28_2315F and myb28_2316R (Tab. S4). Leaves of plants of T_3_ or T_4_ generation were used for GLS analysis to confirm the decrease in aliphatic GLS levels.

### Growth of plant material

All plants were grown in a climatic chamber (PolyKlima, Freising, Germany) at 18°C/22°C, 10h/14h, night/day with 75% light intensity. For propagation, seed production and GLS analysis the plants were sown on regular potting soil (4 L Florador Anzucht soil, 2 L Perligran Premium, 25 g Subtral fertilizer and 2 L tab water) and for amplicon sequencing of naturally colonized leaves, seeds were sown on sieved garden soil from Jena mixed with half the volume of perlite (Perligran Premium). For plant propagation or GLS analyses, unsterilized seeds were sown directly and then vernalized for at least two days at 4°C in the dark. For other experiments, seeds were first surface sterilized using 70% ethanol, followed by 2% bleach and washed 3x with sterile MiliQ water, then were vernalized in 0.1% agarose before sowing. In all cases the plants were thinned out two weeks after germination and if required they were later pricked into individual pots.

### Bacterial growth assays in leaf extract medium

Details on bacterial isolates and how they were recovered from wild plants are found in the supplementary methods and Data S1. We produced plant-based media (“leaf extract medium”) according to (*25*) with minor changes: Leaves of 6-week-old plants (Col-0, *myb28*, *myb28/29,* NG2) were crushed with a metal pestle in R2A broth (1 mL broth for 1 g leaf fresh weight). Supernatants were recovered by centrifugation at maximum speed. Next, the leaf extract media were filter sterilized (0.2 μm) and frozen in aliquots at -80°C. Bacterial isolates were pre-cultured for 24 to 96 h (depending on time required to reach an OD_600_ ≥ 0.2). A 96-well flat-bottom plate was filled with 45 µL leaf extract per well and inoculated with 5 µL of normalized cultures (OD_600_ = 0.2) in triplicates. R2A was added to the negative growth controls. The plate was incubated with constant shaking at 150 rpm at 30 °C. The incubation time reflected the time of the corresponding pre-culture (24-96 h). Directly before the final OD_600_ measurement (VERSAmax^TM^ Microplate Reader, MolecularDevices with SoftMax® Pro Software) 50 µL R2A broth + 0.02% Silwet was added to each well. Raw data from the plate reader was analyzed with custom R scripts. Since growth behavior of strains of the same genus was usually similar, we agglomerated the data on genus level for plotting.

### Evaluation of ITC toxicity on bacterial growth

L-Sulforaphane (4-methyl sulfinyl butyl isothiocyanate, 4MSOB-ITC; ≥ 95 %, CAS 142825-10-3, Sigma-Aldrich) or allyl-ITC (allyl isothiocyanate, AITC; 95 %, CAS 57-06-7, Sigma-Aldrich) were dissolved in DMSO. 3 µL of one ITC or a DMSO control was added to 87 µL R2A medium in 96-well plates resulting in final concentrations ranging from 7.5 to 120 µg/mL. We added 10 μL of each culture normalized to OD_600_ = 0.2 to triplicate wells and covered the plate with a transparent plastic foil to prevent evaporation of the ITCs. The plate was incubated at 30°C in a TECAN Infinite M Plex plate reader and the OD_600_ was measured every 15 min after 1 min of orbital shaking and recorded using the software i-control 2.0. The raw data was processed with custom scripts in R.

### Bacterial growth assays on various aliphatic GLSs as carbon sources

Pre-cultures of individual isolates were washed 2x with 1 mL M9 medium without a carbon source and resuspended in the same medium. The OD_600_ was normalized to 0.2 or 0.3. We dissolved 4MSOB-GLS (glucoraphanin; ≥ 95 %, CAS 142825-10-3, Sigma-Aldrich or Phytoplan Heidelberg, Germany), allyl-GLS (sinigrin; >95 %, CAS 57-06-7, Phytoplan, Heidelberg, Germany) or 2OH3But-GLS (Progoitrin, >97%, CAS 21087-77-4, Phytoplan, Heidelberg, Germany) in sterile MiliQ water to produce 100 mM stocks. M9 minimal medium (12.8 g/L Na_2_HPO_4_, 3.1 g/L KH_2_PO_4_, 0.5 g/L NaCl, 1.0 g/L NH_4_Cl, 0.5 g/L MgSO_4_) (*45*) with final concentrations of 10 mM carbon source or MiliQ water as control was used. Experiments were performed in 96-well plates and the plate was covered with a transparent plastic foil during the incubation time to prevent evaporation. The plate was incubated at room temperature in a TECAN Infinite M Plex plate reader and the OD_600_ was measured every hour after 1 min of orbital shaking. The raw data was processed with custom scripts in R, growth curves were analyzed using the growthcurver package (*71*).

### Enrichment of GLS-utilizing leaf colonizers

To recover isolates which can utilize allyl-GLS as sole carbon source we performed an enrichment similar to (*45*). First, we produced a leaf wash by collecting 10 leaves of 5 NG2 *A. thaliana* plants from the wild NG2 population in spring 2023. For this, leaves were collected and combined in a 1.5 mL tube, they were stored on ice and brought back in the lab. 500 µL 10 mM MgCl_2_ + 0.02% Silwet were added, and the tube was vortexed at lowest possible speed for 20 min. Next, we let the tube stand for ∼5 min to settle down particles and 300 µL of the supernatant was transferred in a new tube. Bacterial load was determined in the leaf wash by plating a 10-fold dilution series on R2A agar. The pH of M9 medium (1.28 g Na_2_HPO_4_, 0.31 g KH_2_PO_4_, 0.05 g NaCl, 0.1 g NH_4_Cl) was adjusted to 7.0 and the medium was autoclaved for 15 min at 121°C. MgSO_4_ stock (5.0 g/L) was prepared separately and filter sterilized. For each enrichment one tube was filled with 70 µL M9 medium, 10 µL MgSO_4_, 10 µL GLS or glucose (100 mM stock) or MiliQ water as control and 10 µL inoculum. Each passage consisted of three replicates with C-source and microbial inoculum and one replicate with water instead of bacteria as control for sterility of the medium. 4MSOB-GLS passages only consisted of one replicate and no water control, due to limited availability of 4MSOB-GLS. Controls with water instead of C-source were inoculated with leaf wash in the beginning, but no growth was observed. The first passage was inoculated with 10 µL of the leaf wash, later passages were inoculated with 10 µL of the previous passage. The tubes were incubated at room temperature in the dark. The incubation time and hence the interval of the passages depended on the time it took the first passage to become visibly turbid (between 3 to 14 days). After each passage, 70 µL were frozen at -80°C with 30 µL 86% sterile glycerol to preserve the microbial communities. A 10-fold dilution series of the replicates of the last passages of allyl-GLS and 4MSOB-GLS enrichments were plated on R2A agar and incubated at 30°C for 48 h. CFUs with different morphologies were picked and isolated. Bacterial strains were identified by sequencing the 16S rRNA gene region with 8F/1492R primers.

### Analysis of GLSs

GLS profiles were measured in leaves and in bacterial cultures. Leaves were collected from lab-grown 4-week-old individuals of our wild isolates from the Jena populations, from 3-week-old plants of T4 generation of NG*myb28* plants to confirm the loss-of-function mutation and wild NG2 and Woe plants. Each genotype was tested at least in three replicates. The plants were harvested by removing roots and flower stems as close to the rosette as possible. The whole rosettes were frozen in liquid nitrogen and kept at -80°C until further processing. GLSs were extracted with methanol and desulfo-(ds)-GLSs were quantified by HPLC coupled to a photodiode array detector as in (*72*). For bacterial cultures, supernatants were recovered by removal of cells after centrifugation at max. speed. Intact GLSs in bacterial cultures were measured directly in the supernatant using HPLC-MS/MS. Details of the analysis procedures are found in the supplementary methods. The following GLSs were detected in the samples: 3-hydroxypropyl GLS (3OHP), 4-hydroxybutyl GLS (4OHB), 3-methylsulfinylpropyl GLS (3MSOP), 4-methylsulfinylbutyl GLS (4MSOB), 2-propenyl GLS (allyl), S-2-hydroxy-3-butenyl GLS (S2OH3But), R-2-hydroxy-3-butenyl GLS (R2OH3But), 3-butenyl GLS (3-Butenyl), 4-pentenyl GLS (4-Pentenyl), 4-hydroxy-indol-3-ylmethyl GLS (4OHI3M), 4-methylthiobutyl GLS (4MTB), 6-methylsulfinylhexyl GLS (6MSOH), 7-methylsulfinylheptyl GLS (7MSOH), 8-methylsulfinyloctyl GLS (8MSOO), indol-3-ylmethyl GLS (I3M), 4-methoxy-indol-3-ylmethyl GLS (4MOI3M), and 1-methoxy-indol-3-ylmethyl GLS (1MOI3M). Results are given as µmol per g dry weight.

### Analysis of GLS breakdown products

GLS breakdown products were measured in leaf homogenates and in bacterial culture supernatants. To analyze GLS breakdown products in leaf homogenates, 120 to 300 mg fresh weight per sample of 6-week-old rosettes of NG2, NG*myb28*, Col-0 and *myb28/29* were harvested. 100 µL MES buffer (50 mM, pH = 6) was added for 100 mg leaf material, next the leaves were grinded with a clean metal pestle and the pestle was rinsed with additional 100 µL MES buffer into the sample. GLS breakdown products were extracted using dichloromethane and measured on a GC-MS and GC-FID. Detected breakdown products were: allyl cyanide (allyl-CN), allyl isothiocyanate (allyl-ITC), 1-cyano-2,3-epithiopropane (CETP), 1-cyano-3,4-epithiobutane (CETB), 4-methylthiobutyl cyanide (4MTB-CN), 4-methylthiobutyl isothiocyanate (4MTB-ITC), 3-methylsulfinylpropyl cyanide (3MSOP-CN), 3-methylsulfinylpropyl isothiocyanate (3MSOP-ITC), 4-methylsulfinylbutyl cyanide (4MSOB-CN), 4-methylsulfinylbutyl isothiocyanate (4MSOB-ITC), 7-methylthioheptyl cyanide (7MTH-CN), 8-methylthiooctyl cyanide (8MTO-CN). Amines in bacterial culture supernatants were measured in the aqueous medium by HPLC-MS/MS. Allyl-ITC and goitrin were extracted from the supernatants with dichloromethane and analyzed by GC-FID. Details of the analysis procedures are found in the supplementary methods.

### 16S rRNA gene amplicon sequencing

The approaches used for amplicon sequencing were slightly different depending on the dataset, an overview is provided in Tab. S5. The three datasets are referred to specifically below.

#### Material and leaf sampling

To analyze bacterial communities in the enrichments in minimal medium (dataset 1, see Tab. S5), the glycerol stocks collected after the passages (see above) were directly used for DNA extraction. To analyze the bacterial community of lab-grown NG2, NG*myb28*, Col-0, *myb28/myb29* leaves (dataset 2, see Tab. S5), 4-6 rosettes of 3-week-old plants per genotype were washed twice with 1 mL sterile MiliQ water by inverting the tube three times to collect “whole” leaves. To collect “endophytes”, an additional 4-6 plants were surface-sterilized with 70% ethanol and 2% bleach (each 1 mL in 1.5 mL tube, 3x inverting) and washed twice with sterile Milli-Q water afterwards. From 2019 to 2020, we collected *A. thaliana* leaf samples from the five different locations in Jena (Tab. S1, dataset 3, see Tab. S5). At the same time, a similar number of other random plants were sampled. Sampling was conducted during the early days of spring in February and March each year. For smaller *A. thaliana* plants, we sampled half of the rosette, while for larger ones, 2-3 leaves were collected. For other plants a similar amount of plant material was selected. The leaf material was washed with sterile MiliQ water three times and samples were brought back to the lab on ice. Plant material was frozen in screw cap tubes with two metal beads and ∼0.2 g glass beads (0.25-0.5 mm diameter) each at -80°C until further processing.

### DNA extraction

Bacterial DNA from GLS enrichments (dataset 1) was extracted from glycerol stocks using an SDS buffer lysis protocol with RNAse A and Proteinase K treatments and a phenol/chloroform cleanup, followed by DNA precipitation. To extract DNA of lab-grown (dataset 2) and wild plants (dataset 3), a CTAB buffer and bead beating protocol was followed, with either a phenol-chloroform cleanup followed by precipitation (dataset 2) or magnetic beads (dataset 3). Precise details of each of the three protocols can be found in the supplementary methods.

### Library preparation for amplicon sequencing

In all library preparations ZymoBIOMICS Microbial Community DNA Standard II (ZymoResearch, Freiburg, Germany; referred to as “ZymoMix”) was used as positive control, and nuclease-free water and CTAB buffer from the DNA extraction were used as negative controls. To quantify bacterial loads in plant samples (dataset 2) we performed a two-step host-associated microbe PCR (hamPCR) with simultaneous amplification of a plant single copy gene (GI = *GIGANTEA* gene, referred to as “GI gene”) along with bacterial 16S rRNA genes according to (*32*). With this protocol, bacterial 16S data can be normalized the GI reads to provide an estimate of bacterial loads in leaves. In all cases, libraries were prepared using a 2-step PCR protocol. In a first 5-cycle PCR, samples were amplified using 341F/799R “universal” 16S rRNA primers modified with an overhang sequence. Plant samples also contained blocking oligos to reduce plastid 16S amplification (*73*) and for dataset 2, GI primers, as mentioned above. The PCR product was enzymatically cleaned to remove remaining primers and used as template in a second, 35-cycle, PCR to add sample index barcodes and sequencing adapters using primers that bound to the overhang region. PCR products were cleaned up with magnetic beads and libraries were quantified using PicoGreen (1:200 diluted stock, Quant-iT^TM^ PicoGreen^TM^, ThermoFisher) in a qPCR machine (qTower^3^, JenaAnalytik, Jena, Germany) or by fluorescence on a gel using ImageJ. Samples were pooled according to their normalized fluorescence relative to the highest fluorescent sample. Pools from the hamPCR protocol with GI were further processed to increase the fraction of 16S relative to GI, as recommended in the original protocol. Libraries were sequenced on an Illumina MiSeq instrument for either 600 cycles (dataset 2,3) or 300 cycles (dataset 1). The precise procedures, including master mix recipes, thermocycling programs and primer sequences can be found in the supplementary materials.

### Data analysis of amplicon sequencing

For all three datasets we first split the amplicon sequencing data on indices and trimmed the adapter sequences from distal read ends using Cutadapt 3.5 (*74*). We then clustered amplicon sequencing data (forward reads only as they were much higher quality) into amplicon sequencing variants (ASVs) using dada2 (*75*). We then removed chimeric sequences and retrieved a sequence table from the merged data. We assigned taxonomy to the final set of ASVs using the Silva 16S rRNA (v 138.1) database (*76*). The database was supplemented by adding the *A. thaliana* GI gene sequence. All positive and negative controls for the datasets were checked. The distribution of taxa in the positive controls were as expected and the negative controls in all cases had <50 reads. In dataset 2 (leaf bacteriomes of lab-grown plants), several Zymomix ASVs (from the positive control) were observed in the negative control and other samples. Since this likely represented low-level background contamination detectable in samples with very low bacterial loads, Zymomix ASVs were removed prior to downstream analysis. We performed downstream analysis in R with Phyloseq (*77*) and VEGAN (*78*) for all three data sets. If applicable, host-derived reads were removed by filtering any ASVs in the order “Chloroplast” and family “Mitochondria” from the 16S ASV tables. For dataset 2, plant GI reads were used to quantify the relative bacterial loads on the plant leaves. To do so, bacterial reads were normalized to the GI reads in each sample. This resulted in small fractions that were not usable with some downstream software, so it was scaled up by multiplying by a factor so that the smallest number in the abundance table is 1. Further analyses for richness, evenness, beta diversity, and differential abundance analysis are described in detail in the supplementary methods.

### Statistical Analysis

All statistical analyses which we performed are mentioned in the respective methods section and in the figure captions in the main text. In addition, raw data (datasets and OTU tables) as well as the R code that generates the figures from the raw data will be publicly available on Figshare before final publication (https://figshare.com/projects/Beyond_defense_Glucosinolate_structural_diversity_shapes_recruitment_of_a_metabolic_network_of_leaf-associated_bacteria/180211).

## Acknowledgements

We acknowledge René Maskos, Stefan Riedel, and Kirsten Küsel (Aquatic Geomicrobiology, Friedrich-Schiller-University Jena) for sequencing our libraries on their MiSeq instrument. Additionally, we acknowledge the help of Beate Rothe in the Biochemistry Department at the Max-Planck-Institute for Chemical Ecology with glucosinolate extractions.

## Funding

Carl Zeiss Foundation via Jena School for Microbial Communication (KU, TM, MTA)

Deutsche Forschungsgemeinschaft (DFG, German Research Foundation) under Germany’s Excellence Strategy - EXC 2051 - Projektnummer 390713860 (TM, MTA)

Deutsche Forschungsgemeinschaft (DFG, German Research Foundation), Projektnummer 458884166 (MTA)

International Max Planck Research School “Chemical Communication in Ecological Systems” (AKR)

Deutsche Forschungsgemeinschaft (DFG, German Research Foundation), Projektnummer 460684957 (UW, AH)

Max Planck Society (JG, MR)

JS received no particular funding for this work.

## Author Contributions

Conceptualization: MTA, KU

Methodology: KU, TM, AKR, MR, AH, JS, MTA

Investigation: KU, TM, AKR, MR, AH, JS

Visualization: KU, AKR

Supervision: MTA, JG, UW

Writing - original draft: MTA, KU

Writing - review & editing: all authors

## Data and materials availability

Data needed to evaluate the conclusions in the paper are present in the paper and/or the Supplementary Materials. Raw sequencing data is available on NCBI-SRA (BioProjects PRJNA1032255 and PRJNA815825) and processed data with code to generate the main figures will be available on Figshare before final publication (https://figshare.com/projects/Beyond_defense_Glucosinolate_structural_diversity_shapes_recruitment_of_a_metabolic_network_of_leaf-associated_bacteria/180211). Local Jena *A. thaliana* genotypes NG2, JT1, SW and Woe are already deposited in NASC database and PB and NG*myb28* mutant will follow.

