## Supplementary Methods and Data for "Beyond defense: Glucosinolate structural diversity shapes recruitment of a metabolic network of leaf-associated bacteria"

**This PDF file includes:**

Supplementary Methods
Figs. S1 to S6
Tables S1 to S7
Data S1 to S2
References (79-93)

**Other Supplementary Materials for this manuscript include the following:**

Data S1 to S2

### Supplementary Methods

*Bacterial isolation, identification and growth conditions*

Details on our microbial collection used in this study are in Data S1. All bacterial isolates used were recovered from *A. thaliana* leaves, the majority from leaves from our local Jena populations. For this, leaves from PB or NG2 population were sampled in spring 2018 or 2019, washed, ground and suspended in PBS + 0.02% Silwet (= leaf extract) to recover microbial leaf colonizers. In both years, we prepared near-extinction dilutions (0 to a few cells per 5 µL) of this leaf extract. In 2018, these dilutions were directly plated on R2A agar or inoculated into R2A broth to isolate bacterial strains. In 2019, the dilutions were inoculated onto axenic germinating Col-0, NG2 and PB seedlings grown on ½ MS + 0.2% sucrose + 1% agar in 24-well plates and two weeks later plants were harvested, crushed and plated to recover only efficient plant colonizers. All bacterial isolates were identified by DNA extraction using an SDS-buffer protocol: 600 µL extraction buffer (0.5 % SDS, 50 mM Tris-HCl pH=8.0, 200 mM NaCl, 2 mM EDTA in NFW) was used to resuspend a cell pellet of an overnight bacterial culture. Samples were bead beaded for 30 s at 1,500 rpm followed by 10 min incubation at 37°C. After centrifugation (20,000 x g, 5 min) the supernatant was recovered. 1/3 vol. 5M potassium acetate were added and centrifuged again, supernatants were cleaned up using 1.5x home-made magnetic Sera Mag purification beads (hereafter "magnetic beads") pre-activated and stored in PEG/NaCl as described in (*79*). PCR amplification of the 16S rRNA genes with 799F/1391R primers (2018 isolates) or 8F/1492R primers (2019 isolates) and Sanger sequencing by Eurofins Genomics (Ebersberg, Germany) was performed (all primers are listed in Tab. S4). All isolates were cultured in R2A broth at 30°C, 150 rpm, if not stated otherwise.

*Whole genome sequencing of bacterial strains*

Bacterial DNA was extracted using the same SDS buffer protocol as described above. The samples were treated with RNAse A (2 µg/mL, 37°C for 30 min), followed by a treatment with Proteinase K (100 µg/mL, 37°C for 1 h). We then purified the samples twice by adding an equal volume of phenol/chloroform/isoamyl alcohol and recovering the aqueous layer. The DNA was precipitated by adding sodium acetate and ethanol overnight. In total we sent 30-40 µL with a DNA concentration of at least 50 ng/µL for whole genomes sequencing. The genomes were sequenced on a NextSeq 2000 platform (Illumina Sequencing) at a depth of 300 MBp at Microbial Genome Sequencing Center (Pittsburgh, PA 15222, USA). The results were provided as paired end reads

(2x151bp, fastq files). We assembled and annotated the sequences using an in-house perl script. We trimmed the raw reads with the software TrimmomaticPE (*80*) and then assembled them with the software SPAdes Version 3.14.1 (*81*) using default k-mer sizes of 21, 33, 55 and 77. The resulting scaffolds fasta file was then annotated with PROKKA 1.14.6 (*82*) and the reference genome Pseudomonas_syringae_pv_tomato_DC3000_111.gbk retrieved from NCBI database.

*Analysis of 4MSOB-ITC and 4MSOB-amine in bacterial cultures by LC-MS/MS*

4MSOB-ITC in bacterial cultures was analyzed on an Agilent 1200 HPLC system (Agilent, Santa Clara, CA, United States) coupled to an API3200 tandem mass spectrometer (AB SCIEX, Darmstadt, Germany). Amines were separated by an Agilent XDB-C18 column (5 cm × 4.6 mm, 1.8 µm, Agilent, Waldbronn, Germany). The mobile phase consisted of 0.05% (v/v) formic acid in ultrapure water as solvent A and acetonitrile as solvent B, at a flow rate of 1.1 mL/min. The elution gradient was: 0-0.5 min, 3-15% B; 0.5-2.5 min, 15-85% B; 2.5-2.52 min, 85-100% B; 2.25-3.5 min, 100% B; 3.5-3.51 min, 100-3% B, 3.51-6 min, 3% B. The ion spray voltage was maintained at 5500 eV in positive mode. The turbo gas temperature was set to 500 °C, nebulizing gas to 60 psi, drying gas to 60 psi, curtain gas to 35 psi, and collision gas to 3 psi. Details of multiple reaction monitoring (MRM) can be found in Tab. S7. Analyst Software 1.6 Build 3773 (AB SCIEX) was used to acquire and process data.

*Detailed analysis of GLSs in leaves and bacterial cultures*

For leaf material, the measurement was performed as previously described by (*83*). After freeze-drying, rosettes were ground to powder by shaking with metal beads. GLSs were extracted from 8-12 mg of the powder or 1-2 mg in case of axenic seedling by 1 mL of 80 % MeOH with 50 µM pOH-Benzyl GLS (Sinalbin, internal standard) and thorough vortexing for 30 sec and shaking for 30 min at high speed. The samples were centrifuged for 5 min at 3,200 rpm and 600 µL or 800 µL of the supernatant was added to a freshly prepared DEAE-Sephadex-filter plate and allowed to flow through. Samples on the DEAE-Sephadex were washed five times with: (1) 0.5 mL 80% MeOH, (2/3) two times 1 mL ultrapure water, (4) 0.5 mL 0.02 MES pH 5.2 and (5) 30 µL sulfatase solution. Samples were kept at room temperature overnight and desulfo- (ds)-GLSs were eluted with 0.5 mL ddwater on the next day. The eluted ds-GLSs were separated using high performance liquid chromatography (Agilent 1100 HPLC system, Agilent Technologies) on a reversed phase column (Nucleodur Sphinx RP, 250 x 4.6 mm, 5µm, Macherey-Nagel, Düren, Germany) with a water (A)-acetonitrile (B) gradient (0-1 min, 1.5% B; 1-6 min, 1.5-5% B; 6-8 min, 5-7% B; 8-18 min, 7-21% B; 18-23 min, 21-29% B; 23-23.1 min, 29-100% B; 23.1-24min 100% B and 24.1-28 min 1.5% B; flow 1.0 mL min-1). Detection was performed with a photodiode array detector and peaks were integrated at 229 nm. We used the following response factors: 3OHP and 4OHB 2.8, all other aliphatic GLS 2.0, indole GLS 0.5 (*72*) for quantification of individual GLSs. The identity of the peaks was based on a comparison of retention time and UV absorption spectrum with data obtained for isolated ds-GLSs as described in (*84*) and by analysis of the ds-GLS extracts on an LC-ESI-Ion-Trap-mass spectrometer (Esquire6000, Bruker Daltonics).

For bacterial culture supernatants, after centrifugation at max. speed the supernatants were recovered. The levels of GLSs were quantified in 1:10 (v:v) diluted extracts using an Agilent 1200 HPLC system (Agilent, Santa Clara, CA, United States) coupled to an API3200 tandem mass spectrometer (AB SCIEX, Darmstadt, Germany) according to (*85*). GLSs were separated on an EC 250/4.6 NUCLEODUR Sphinx RP column (250 mm × 4.6 mm, 5 µm; Macherey-Nagel, Düren, Germany). The mobile phase consisted of 0.2% (v/v) formic acid in ultrapure water as solvent A and acetonitrile as solvent B, at a flow rate of 1 mL/min. The elution gradient was: 0-1 min, 1.5% B; 1-6 min, 1.5-5% B; 6-8 min, 5-7% B; 8-18 min, 7-21% B; 18-23 min, 21-29% B; 23-23.1 min, 29-100% B; 23.1-24 min, 100% B; 24-24.1 min, 100-1.5% B; 24.1-28 min, 1.5% B. The ionization source was set to negative mode. The ion spray voltage was maintained at -4,500 eV. Gas temperature was set to 500 °C, nebulizing gas to 60 psi, drying gas to 60 psi, curtain gas to 30 psi and collision gas to 6 psi. For each compound, the transitions from precursor ion to product ion was monitored using multiple reaction monitoring (MRM) (Tab. S6). Compounds were quantified using external calibration curves generated using the following compounds: Allyl-GLS (Carl Roth, Mannheim, Germany), 4-methylsulfinylbutyl-GLS (4MSOB-GLS, Phytoplan, Heidelberg, Germany), and 2OH3But-GLS (Progoitrin, Phytoplan, Heidelberg, Germany). Analyst Software 1.6 Build 3773 (AB SCIEX) was used to acquire and process data.

*Detailed analysis of GLS breakdown products in leaf homogenates and bacterial cultures*

For leaf homogenates, internal standards (25 µL of 100 ng/µL phenyl cyanide in methanol, 20 µL of 1 nmol/µL indole-3-carbonitrile in methanol) were added to each sample after crushing the leaf material in MES buffer. After 10 min total incubation time, the samples were centrifuged at 13,400 rpm at 20°C for 10 min. The supernatant was transferred into 1.5 mL glass vials and frozen at -20°C until further processing. Thawed supernatants were extracted twice with 750 µL dichloromethane. The organic phases were pooled and dried over $Na_2SO_4$, concentrated in an air stream to about 150 µL and analyzed by GC-MS and GC-FID for aliphatic hydrolysis products as described in (*86*), using a ZB-5MS capillary column (30 m × 0.25 mm; ft 0.25 µm; Phenomenex, Aschaffenburg, Germany).

To analyze GLS breakdown product generated by R3, 200 µL cultures in M9 medium with 10 mM allyl-GLS were grown for 150 h and with 10 mM 2OH3But-GLS for 220 h in a 96-well plate as described above. 10 µL of the supernatants were diluted 1:10 in ddwater to measure amines after pre-tests had shown too high peaks of undiluted samples. Amines were analyzed on an Agilent 1200 HPLC system (Agilent, Santa Clara, CA, United States) coupled to an API3200 tandem mass spectrometer (AB SCIEX, Darmstadt, Germany). Amines were separated by an Agilent XDB-C18 column (5 cm × 4.6 mm, 1.8 µm, Agilent, Waldbronn, Germany). The mobile phase consisted of 0.05% (v/v) formic acid in ultrapure water as solvent A and acetonitrile as solvent B, at a flow rate of 1.1 mL/min. The elution gradient was: 0-0.5 min, 3-15% B; 0.5-2.5 min, 15-85% B; 2.5-2.52 min, 85-100% B; 2.25-3.5 min, 100% B; 3.5-3.51 min, 100-3% B, 3.51-6 min, 3% B. The ion spray voltage was maintained at 5500 eV in positive mode. The turbo gas temperature was set to 500 °C, nebulizing gas to 60 psi, drying gas to 60 psi, curtain gas to 35 psi, and collision gas to 3 psi. For each compound, the transitions from precursor ion to product ion was monitored using multiple reaction monitoring (MRM) (Tab. S7). Compounds were quantified using an external calibration curve generated with allyl-amine (Sigma-Aldrich, Taufkirchen, Germany). Analyst Software 1.6 Build 3773 (AB SCIEX) was used to acquire and process data.

10 µL of Phenylcyanide (1:10000 dilution in MeOH; Merck, Darmstadt, Germany) was added as internal standard to 140 µL of the supernatant. Nonpolar compounds were extracted with 400 µL dichloromethane, vortexing and centrifugation for 1 min at 3,500 x g. About 100 µL of the organic phase was transferred into fresh glass vials. Allyl-ITC and goitrin were quantified following the method described in (*87*). Samples were analyzed by GC-FID using an Agilent 6890 Series gas chromatograph with an DB5MS column (30 m x 0.25 mm x 0.25 m film, Agilent Technologies), splitless injection at 200°C, and a temperature program of 35°C for 3 min, a 12°C/min ramp to 96°C and a 18°C/min ramp to 240°C (with a 6 min final hold). Peaks were identified by comparison of retention times and mass spectra to those of the authentic standards, allyl-ITC

(Fluka), and goitrin (Sigma-Aldrich). Hydrogen was used as the carrier gas and the detector operated at 300°C. Quantification was based on peak area relative to that of the internal standard, phenylcyanide. Response factors (RF) relative to phenylcyanide were experimentally determined for allyl-ITC (1.55) and calculated for goitrin (1.79) by the effective carbon number (ECN) concept (*88*).

*DNA extraction from enrichment cultures (dataset 1)*

Bacterial DNA from GLS enrichments (dataset 1, Tab. S5) was extracted from glycerol stocks using the same SDS buffer protocol as described earlier for individual bacterial strains. To recover higher quality DNA the resulting supernatants were cleaned up using RNAse A (10 µg/mL, 30 min at 37°C) and proteinase K treatment (100 µg/mL, 1 h at 37°C), followed by a two-step purification: equal amounts of phenol/chloroform were added to the supernatants, followed by centrifugation at 20,000 x g for 10 min at 4°C and recovery of the aqueous phase. Next, equal amounts of chloroform/isoamyl alcohol (24:1) were added, followed by centrifugation at 20,000 x g for 10 min at 4°C and recovery of aqueous phase. The DNA was precipitated by adding 1/10 Vol. sodium acetate (3M, pH5) and 2.5 Vol. isopropanol, incubation at -20°C overnight and 30 min centrifugation at 4°C at 20,000 x g. The DNA pellet was washed twice with 80% ethanol and air-dried before resuspending in 10 mM Tris-HCl pH8.

*DNA extraction from lab and wild plants (datasets 2 and 3)*

To extract DNA from plant material of wild *A. thaliana* plants for dataset 3 (Tab. S5), all tubes with leaf material were bead beat for 30 s at 1,400 rpm. 150 µL of 2x CTAB buffer (2% cetyltrimethylammonium bromide, 1% polyvinylpyrrolidone, 100 mM Tris-HCl, 1.4 M NaCl, 20 mM EDTA) was added to each tube, vortexed briefly and incubated at 37°C for 10 min. The tubes were spun down for 5 min at 20,000 x g and the supernatant was transferred into a deep 96-well plate. 1/10 Vol. of sodium acetate (3M, pH5) and 2.5 Vol. of 100% ethanol were added to each sample and the plate was frozen at -20°C to precipitate DNA overnight. To collect the DNA the plate was centrifuged at 20,000 x g for 60 min at 4°C. All supernatants were discarded, and the pellets were washed twice with 70% ethanol, airdried and re-suspended in 50 µL 10 mM Tris-HCl (pH8). The samples were frozen at -20°C overnight and cleaned up using 1.5x magnetic beads as described earlier. To extract DNA of lab-grown plants (dataset 2), all tubes with leaf material were bead beat for 30 s at 1,400 rpm. 300 µL of 2x CTAB buffer was added to each tube, vortexed briefly and incubated at 65°C for 15 min. The tubes were spun down for 5 min at 20,000 x g and the supernatant was transferred into a fresh tube. Equal amounts of Phenol:Chloroform:Isoamyl alcohol (25:24:1) were added, briefly vortexed and centrifuged for another 5 min at 20,000 x g. The DNA was precipitated and eluted as described for data set 3, without additional magnetic bead clean-up.

*Detailed amplicon sequencing procedure*

Master mixes for the first PCR for dataset 2 and 3 contained 11.36 µL nuclease-free water, 4.00 µL 5x Kapa High Fidelity Buffer, 0.60 µL 10 mM Kapa dNTPs, 0.16 µL of each primer (GI_F, GI_R, 341F-OH, 799R-OH, Tab. S4), 0.50 µL of each blocking Oligo (At_BLc_16S_F5, At_BLc_16S_R1, Tab. S4), 0.40 µL Kapa HiFi polymerase (Kapa Biosystems), 2.00 µL normalized DNA (50-100 ng/µL) as template per reaction. Blocking oligos bound to their targets in the first PCR in a BioRad thermocycler: (1) 95°C for 3:00 min, (2) 98°C for 0:20 min, (3) 58°C for 0:30 min, (4) 55°C for 1:00 min, (5) 72°C for 1:00 min, to step (2) and repeat for 5 cycles, (6) 72°C for 2:00. All samples were cleaned up enzymatically by adding 0.50 µL Exonuclease I, 0.50 µL Antarctic Phosphatase and 1.22 µL Antarctic Phosphatase Buffer (NewEngland Biolabs) to 10 µL of the first PCR reaction at 37°C for 30 min, followed by enzyme deactivation at 80°C for 15 min. In the second PCR the master mix consisted of 7.34 µL nuclease-free water, 4.00 µL Kapa High Fidelity Buffer, 0.6 µL 10 mM dNTPs, 0.40 µL Kapa polymerase, 5 µL of cleaned first PCR reaction, along with unique forward and reverse indexing primers for each sample. Products were amplified: (1) 95°C for 3:00 min, (2) 98°C for 0:20 min, (3) 60°C for 1:00 min, (4) 72°C for 1:00 min, to step (2) and repeat for 35 cycles, (5) 72°C for 2:00. All samples were cleaned up using 1.5x vol. magnetic beads as described earlier in the methods.

To pool the libraries of datasets 2 and 3, the fluorescence of each sample was measured with Picogreen (1:200 diluted stock, Quant-iT™ PicoGreen™, ThermoFisher) in a qPCR machine (qTower$^3$, JenaAnalytik, Jena, Germany). Samples were pooled according to their normalized fluorescence relative to the highest fluorescent well. Dataset 2 contained plant GI reads and the entire volume of the final library ran on a 2 % high resolution agarose gel to separate GI and 16S bands. Both were purified from the gel using the GeneJET gel purification kit (ThermoFisher). Additionally, we re-barcoded the entire library using 1-5 ng of the pooled library according to (*32*) with minor modifications. The PCR master mix contained 7.34 µL nuclease-free water, 4.00 µL 5x Kapa Buffer, 0.60 µL 10 mM Kapa dNTPs, 0.40 µL Kapa enzyme, 1.33 µL of 4.5 µM of forward and reverse indexing primers and 5 µL of the final pool. Products were amplified in 8 cycles with the protocol for the second PCR mentioned above for dataset 2. Afterwards the re-barcoded library was cleaned up using 1.5x magnetic beads. In the end, the fluorescence of the two fractions (GI, 16S) along with the re-barcoded library was determined and the fractions were combined (94% 16S rRNA, 5% GI, 1% re-barcoded library).

For bacterial-only communities from GLS enrichments (dataset 1) the PCR protocols were the same as mentioned above with minor exceptions. As there is no need to block plant 16S reads or quantify GI reads we only used 341F-OH/799R-OH primers for amplification, instead the volume of DNA template was increased to 2.5 µL per 10 µL reaction. Because of unspecific bands after the second PCR, pooling was performed according to the brightness of the 16S rRNA gene bands (660 bp) on a gel using ImageJ. The samples were normalized relative to the brightest sample and pooled into sub-pools. Libraries were sequenced on a Illumina MiSeq instrument for either 600 cycles (dataset 2,3) or 300 cycles (dataset 1).

*Analysis of diversity of bacterial communities based on 16S rRNA gene data*

Richness and evenness (Chao1, Shannon, Simpson, ACE) in datasets 1 and 2. To plot bar charts and calculate beta diversity matrices we excluded samples with less than 100 bacterial reads in all three datasets. For the bacterial communities in the enrichments (dataset 1), we agglomerated the taxa at genus level and taxa with less than 0.02 % abundance in one sample were agglomerated and classified as “Remainder” in the bar charts. For dataset 1 we used relative abundance data to calculate beta diversity distances, and dataset 2 was normalized to plant GI reads. In both cases, beta diversity is based on the Bray-Curtis and Jaccard distance and ordination was performed with principle coordinate analysis (PCoA). For visualization purposes we constrained by genotype. The statistical significance of explaining variables was tested with a permutational analysis of variance (PERMANOVA) test. For dataset 3, we eliminated samples with fewer than three biological replicates per treatment and agglomerated on genus level. Beta diversity was calculated as the Aitchison distance between samples, which applies a centered log ratio (CLR) transformation followed by calculating Euclidean distances. The resultant patterns were illustrated in ordination plots that were faceted by year and plant type to provide a comprehensive overview. The influence of various factors, including plant type, location, year, and month, as well as the interaction of the variables on microbial community structures was assessed using PERMANOVA (significance level $p<0.05$).

To perform differential abundance analysis (DESeq2 (*89*)) on pairwise comparisons, we added a pseudo count of 1 to each read in the ASV table of non-normalized counts in dataset 3 or plant GI-normalized reads in dataset 2. Size factors were estimated based on the geometric means of each taxon. A Wald test with a parametric fit was executed to determine significant differences. To control the false discovery rate, the Benjamini-Hochberg method was adopted. Taxa with adjusted p-values below 0.05 were marked as significantly differentially abundant. log10-transformed abundances were plotted and a Wilcox test was applied to confirm the significance.

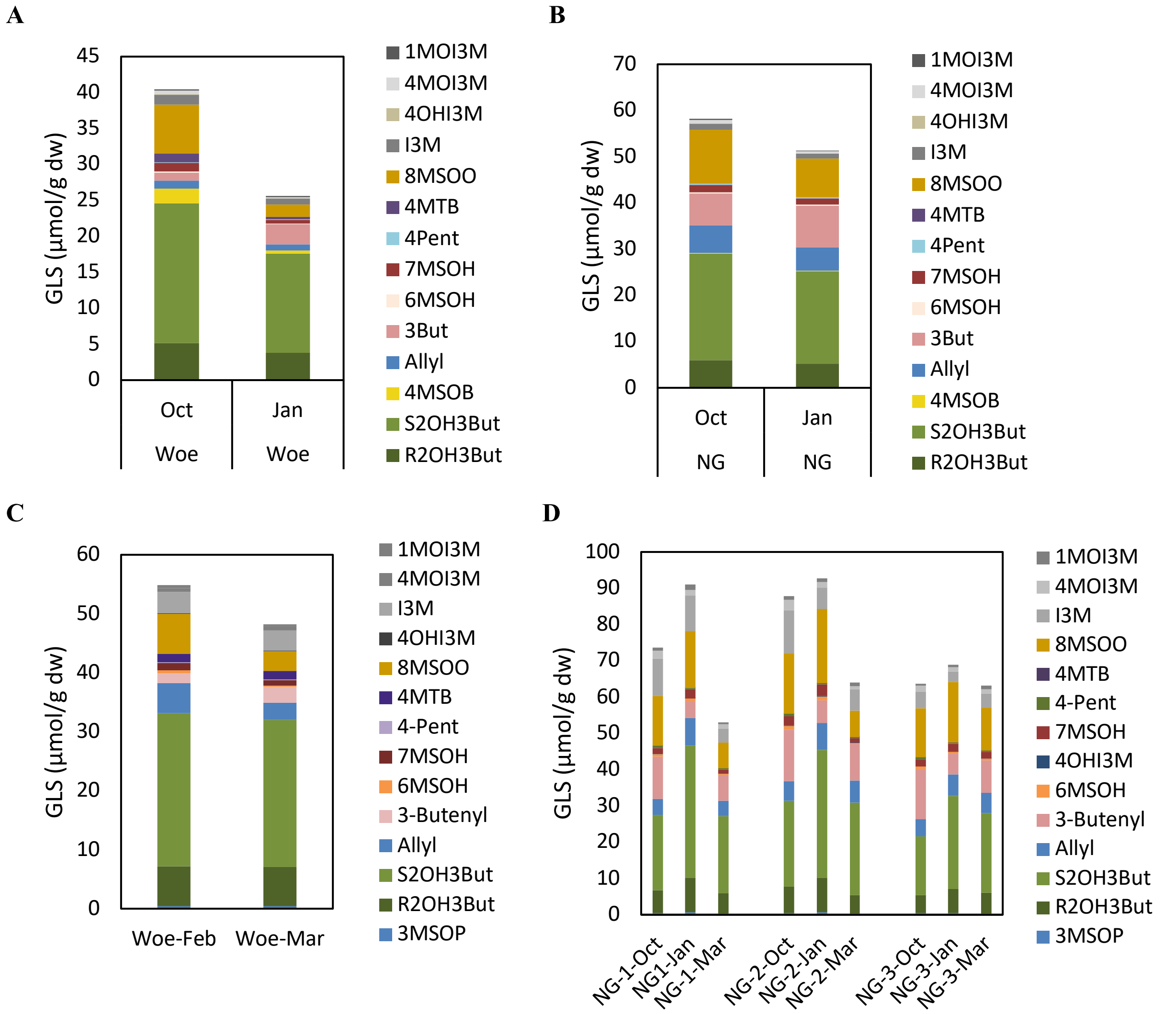


**Fig. S1.**

**GLS profiles of wild *A. thaliana* populations NG2 and Woe.** Average GLS concentration of 5-6 replicates sampled at Woe (**A**) and NG2 (**B**) in October 2021 and January 2022. (**C**) Average GLS concentration of 5-6 replicates of Woe sampled in February and March 2023. (**D**) Average GLS concentration of 5 replicates of three sub-plots at NG2 location sampled in October 2022, January and March 2023. Abbreviations of GLSs are listed in the methods section.

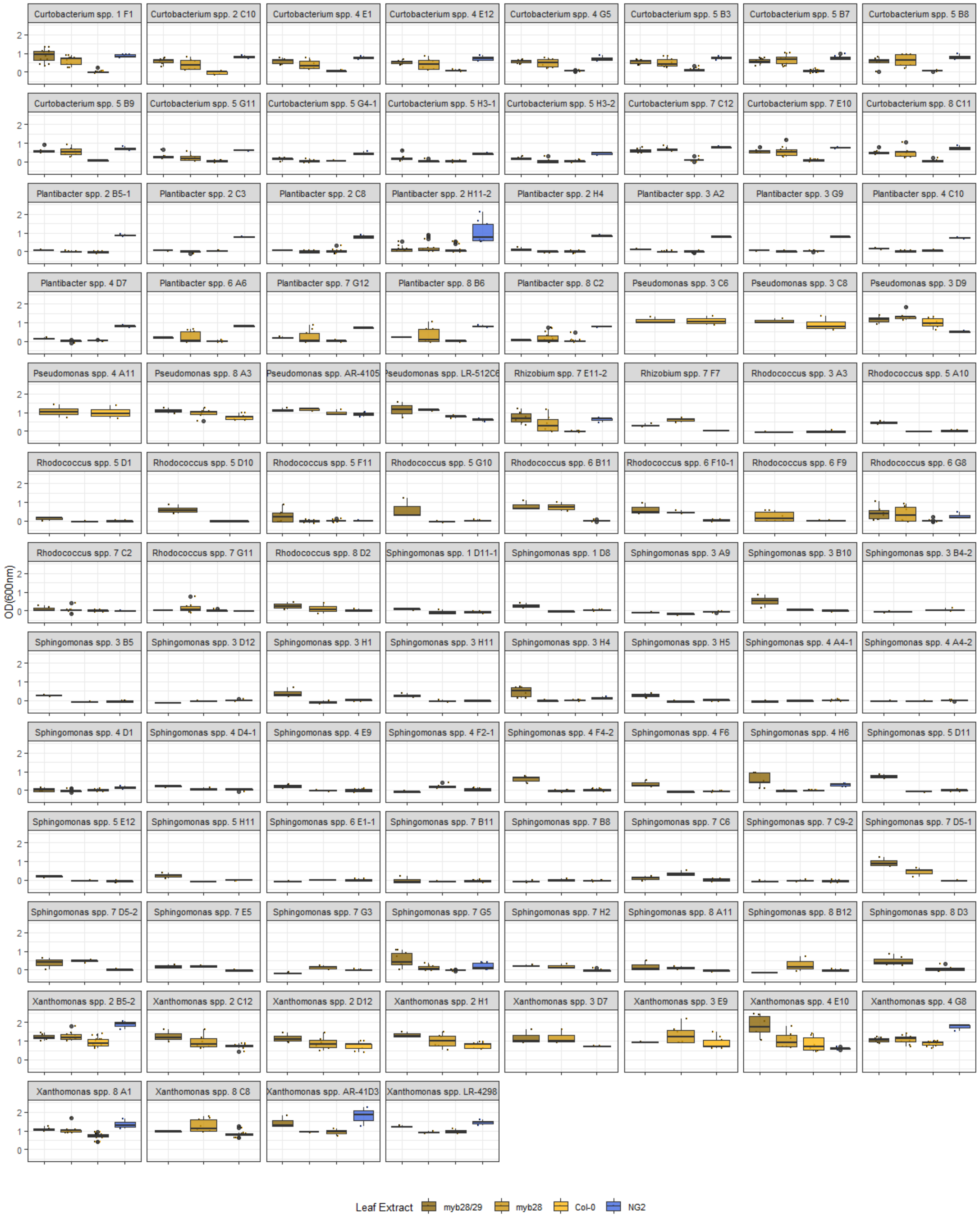


**Fig. S2.**

**Bacterial growth in leaf extract medium after 24-96 h incubation time.** The plots represent individual results for all strains which are shown on genus level in Fig. 2A.

**A**

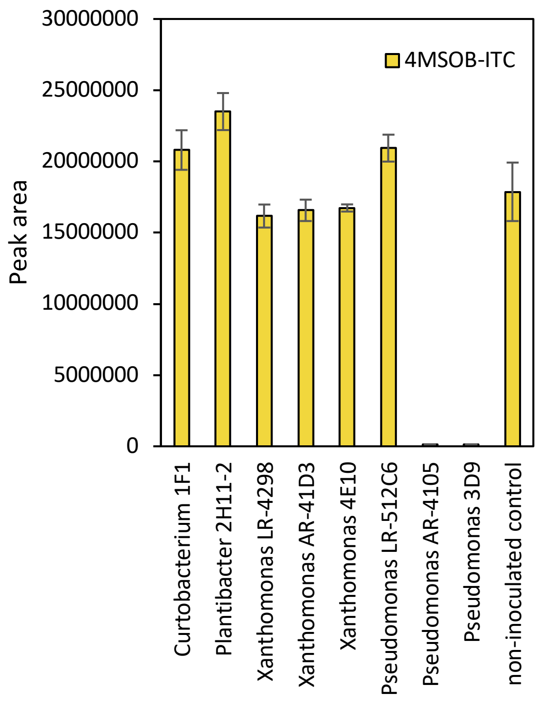


**B**

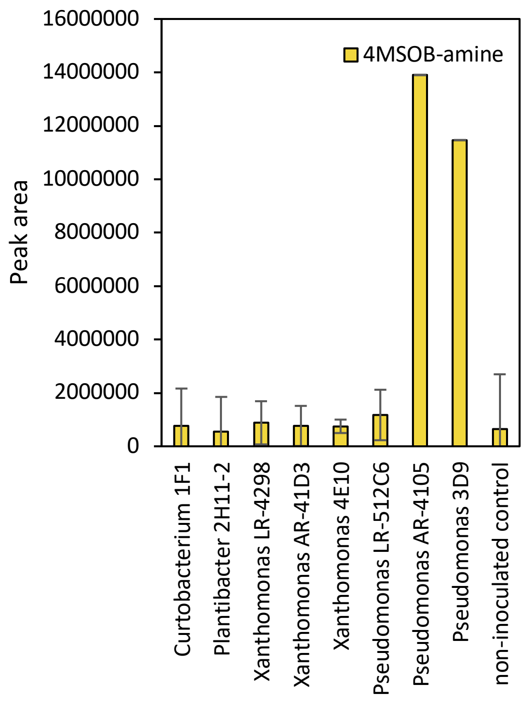


**Fig. S3.**

**Bacterial 4MSOB-ITC degradation *in-vitro*.** Bacterial strains were grown for 18 h in R2A broth supplemented with 30 µg/mL 4MSOB-ITC or DMSO as control. 4MSOB-ITC and its *saxA*-mediated breakdown product 4MSOB-amine were measured in the aqueous supernatants without further extraction. (**A**) average peak area of 4MSOB-ITC from LC-MS/MS of three replicates per strain (**B**) average peak area of 4SMBO-amine from LC-MS/MS of the same three replicates per strain.

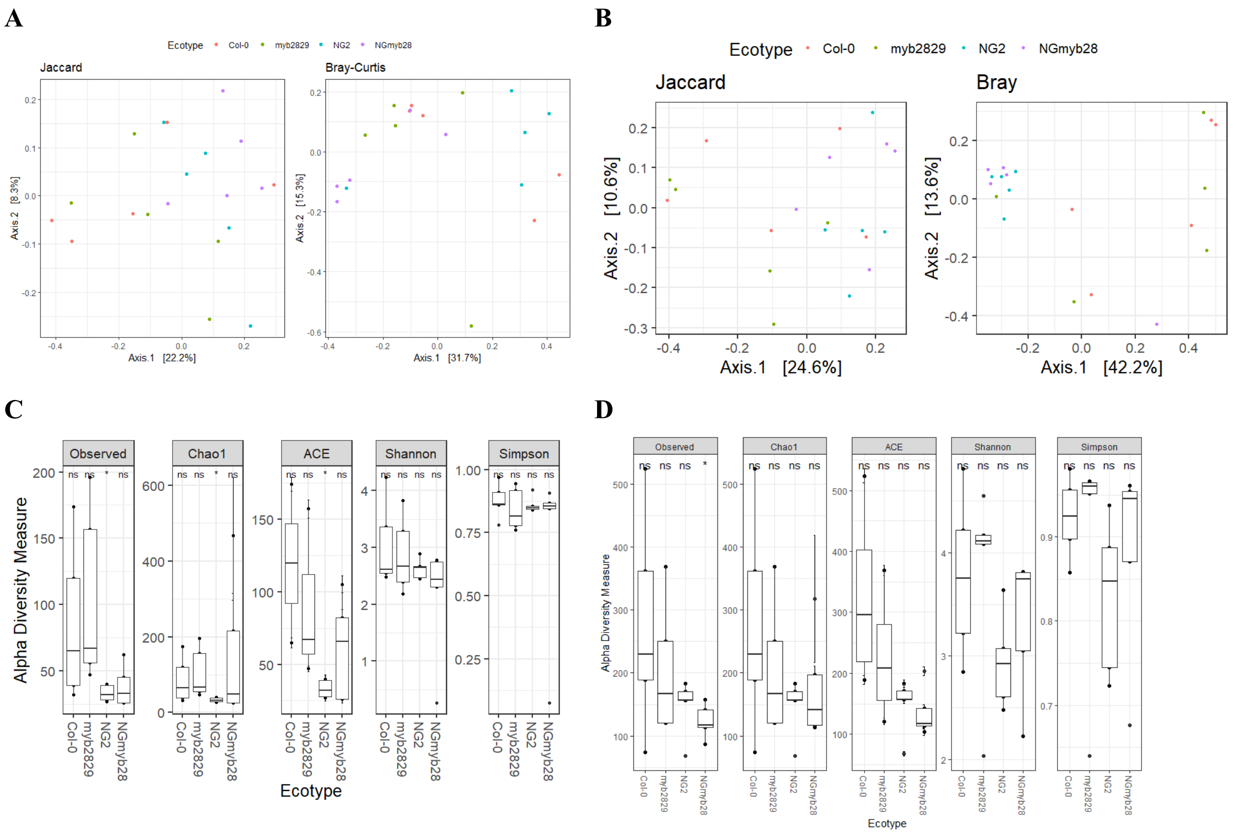


**Fig. S4.**

**Alpha and beta diversity of leaf bacterial communities of lab-grown NG2, NG*myb28*, Col-0 and *myb28/myb29* plants.** Plants were grown for 3-4 weeks and bacterial communities were assessed in either surface sterilized (**A**,**C**) or washed leaves (**B**,**D**). 5-6 replicates per genotype with >100 reads per sample were included. To calculate beta diversities, the data was agglomerated on genus level, and reads were normalized to the plant GI reads. Detailed statistical analyses for all plots are presented in Supplementary Data S2 (**A**) unconstraind PCoA plots of beta diversity (Jaccard, Bray-Curtis) in endophytic community data. (**B**) Unconstrained PCoA plots of beta diversity (Jaccard, Bray-Curtis) in total community data. (**C**) Alpha diversity of non-normalized samples for endophytic communities. (**D**) Alpha diversity of non-normalized samples for total communities.

**A**

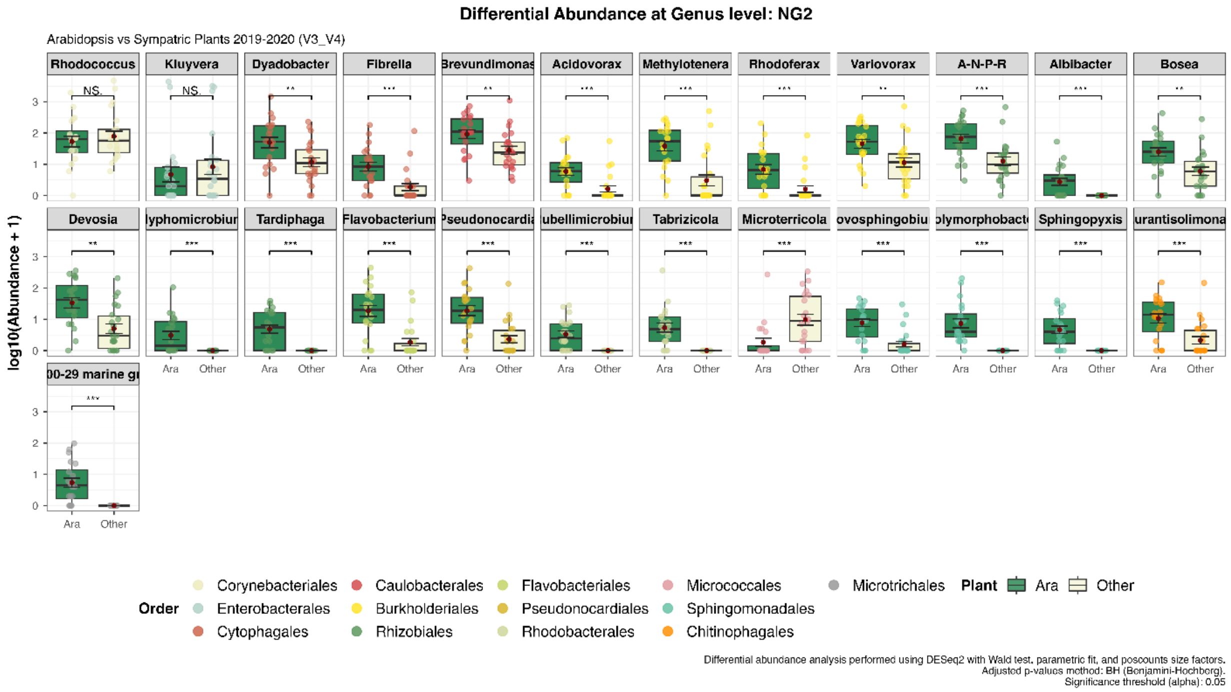


**B**

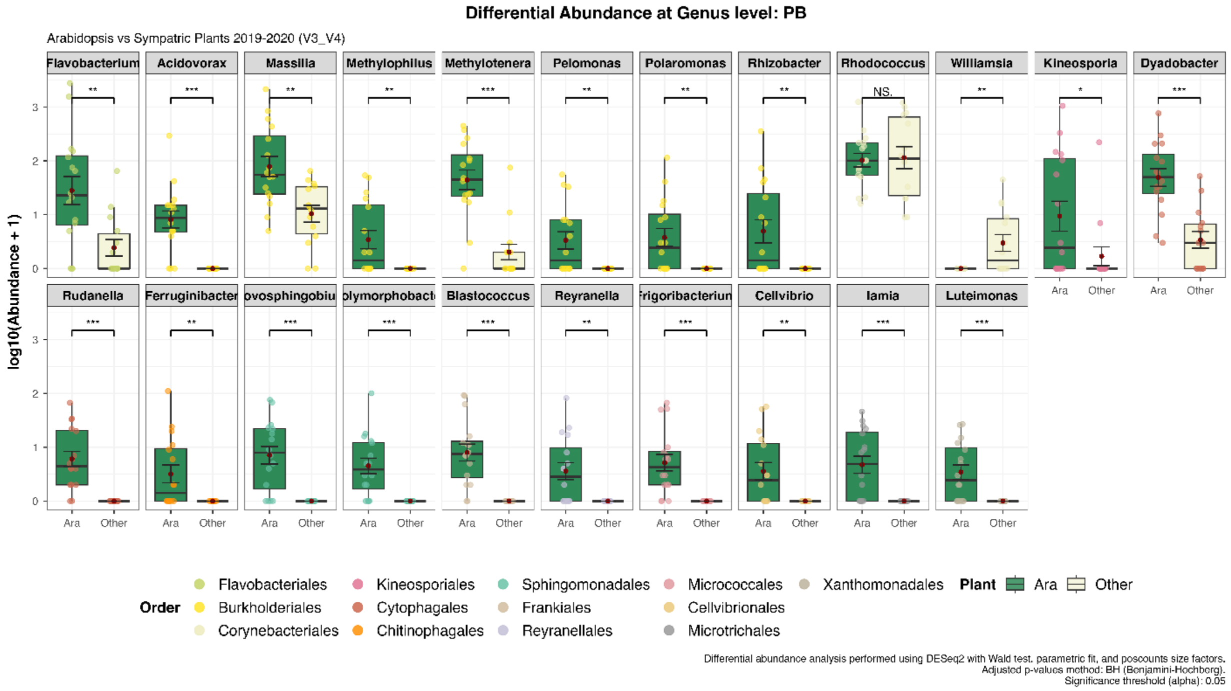


*The figure is ongoing on the next page.*

*The figure is ongoing from the previous page.*

**C**

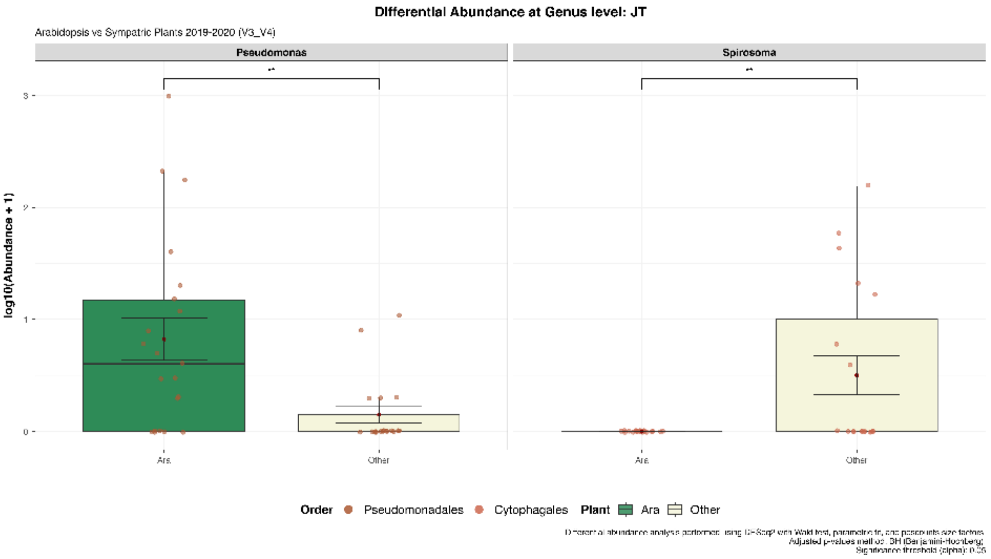


**D**

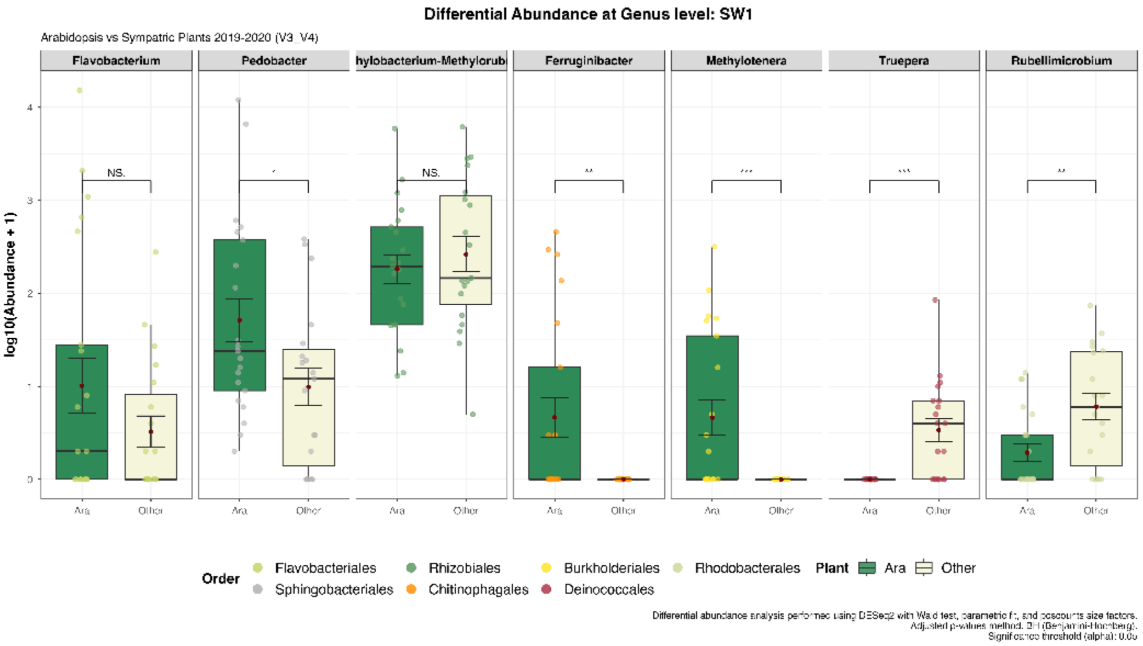


*The figure is ongoing on the next page.*

*The figure is ongoing from the previous page.*

**E**

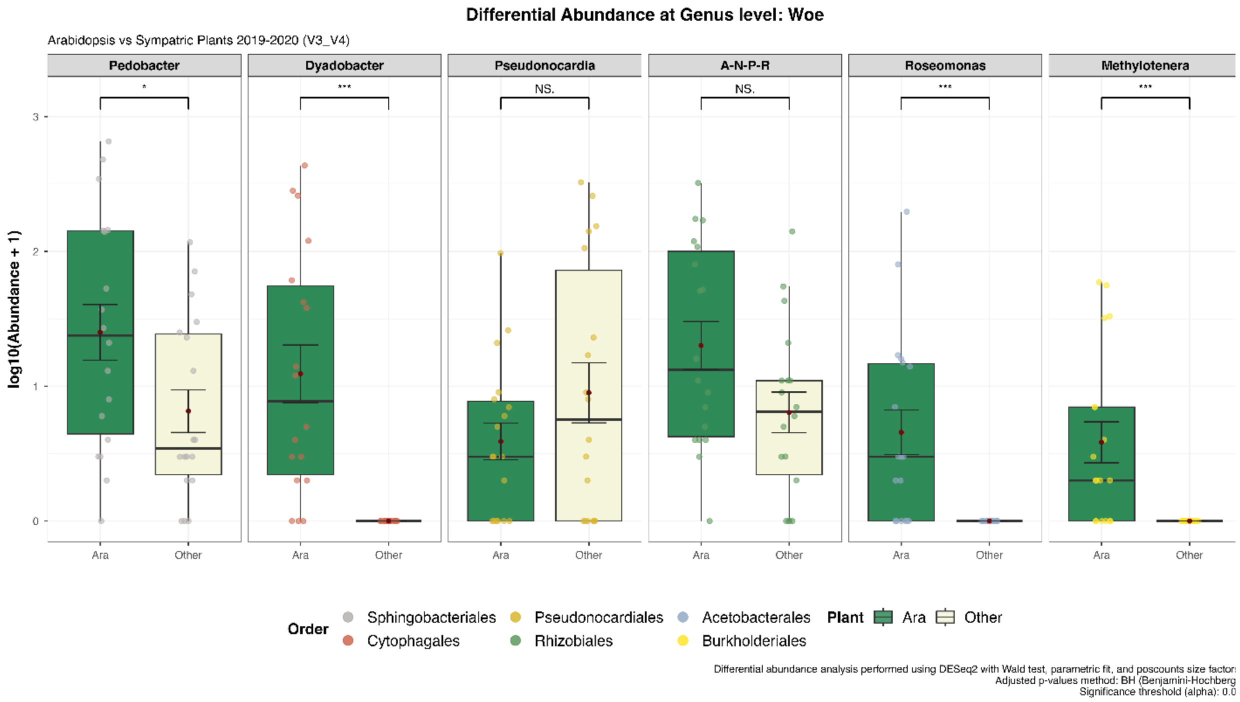


**Fig. S5.**

**Differential abundant taxa in leaf bacterial communities from *A. thaliana* compared to sympatric plants across the five Jena locations in February and March 2019 and 2020.** Samplings with < 3 replicates were excluded and low abundance taxa were filtered out. The pseudocount of 1 was added to each sample prior to DESeq analysis (cutoff alpha = 0.05). All significantly different taxa are plotted with p-values referring to differences in abundance based on Benjamini-Hochberg adjustment. The five locations were visualized separately (**A**) NG2, (**B**) PB, (**C**) JT1, (**D**) SW1 and (**E**) Woe.

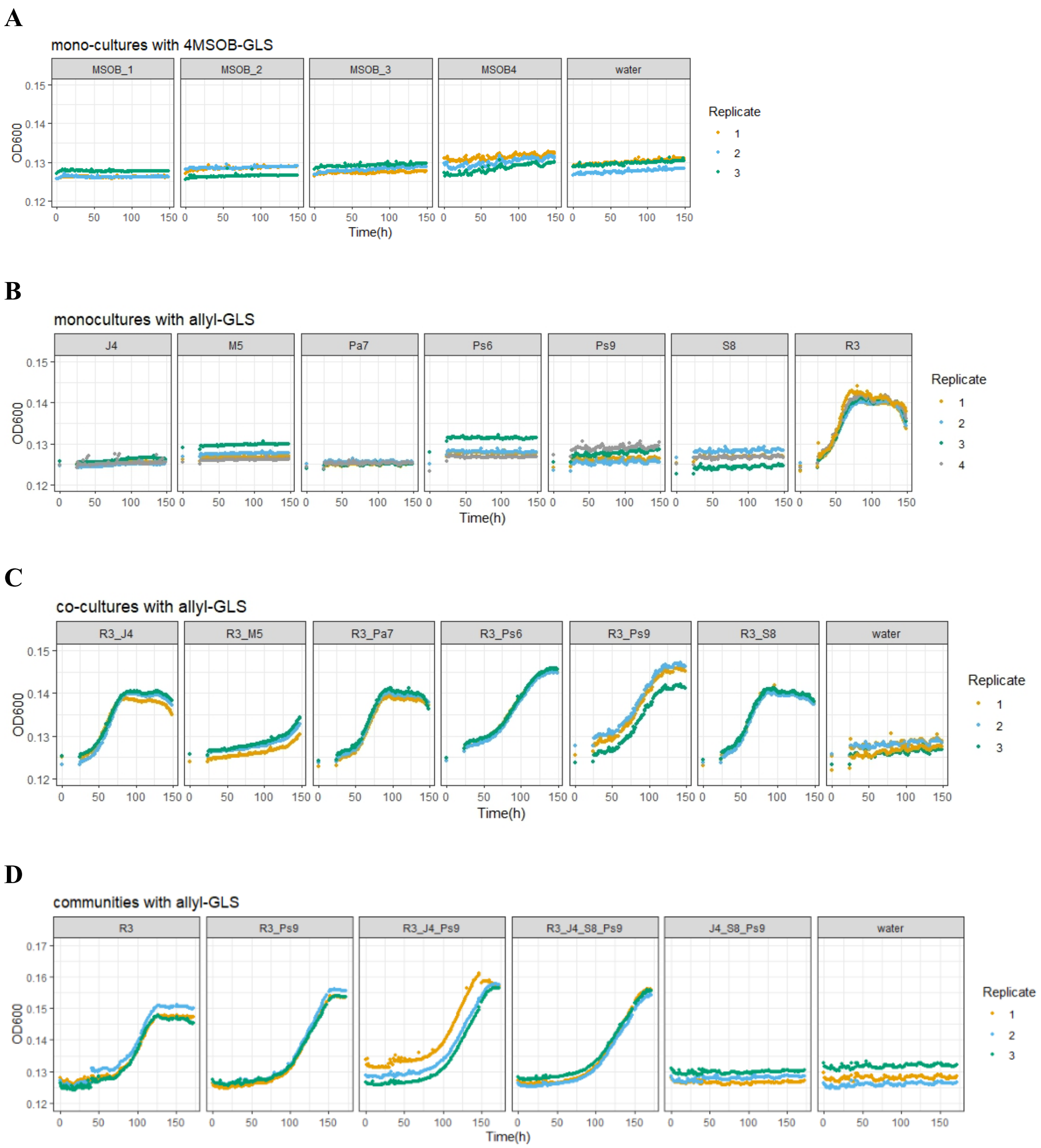


**Fig. S6.**

**Growth of bacterial isolates on glucose and aliphatic GLSs**. All isolates were grown in M9 medium supplemented with 10 mM C-source in one six-day passage. $OD_{600}$ was measured every hour, n=3-4, water served as negative control. (**A**) Monocultures of *Pseudomonas* strains with 4MSOB-GLS from leaf enrichment in M9 with 4MSOB-GLS. (**B**) Monocultures in media supplemented with allyl-GLS of all seven isolates which were recovered from allyl-GLS enrichments, data of the first 24 h is missing. (**C**) Co-cultures of the same isolates shown in C with R3 with allyl-GLS, data of the first 24 h is missing. (**D**) Communities of R3 with different strains with allyl-GLS.

**Table S1.**

**The five local *A. thaliana* populations identified in the city of Jena, Germany.**

| Name | Date found | Date collected | Name in NASC database |
|---|---|---|---|
| NG2 | Fall 2017 | 09.05.2018 | Je-1 |
| PB | Fall 2017 | 13.03.2018 | Not deposited yet |
| Woe | 05.04.2018 | 05.04.2018 | Je-3 |
| SW1 | 12.04.2018 | 17.04.2018 | Je-4 |
| JT1 | 17.04.2018 | 17.04.2018 | Je-2 |

**Table S2.**

***sax* gene content in draft whole genomes of leaf colonizers.** Annotations are based on the genome of *Pseudomonas syringae* pv. tomato DC3000. Number of X represent the number of gene copies.

| ID | Isolate | *saxA* | *saxC* | *saxB* | *saxD* | *saxF* | *saxG* |
|---|---|---|---|---|---|---|---|
| 5B7 | *Curtobacterium* sp. | | | | | | |
| F1 | *Curtobacterium* sp. | | | | | | |
| 2H11-2 | *Plantibacter* sp. | | | | | | |
| LR-4298 | *Xanthomonas campestris* | | | X | XXX | | |
| AR-41D3 | *Xanthomonas campestris* | | | X | XXX | | |
| 4E10 | *Xanthomonas campestris* | | | | XXX | X | |
| LR-512C6 | *Pseudomonas koreensis* | | | X | XX | X | X |
| AR-4105 | *Pseudomonas syringae* | X | X | X | XXXX | X | X |
| 3D9 | *Pseudomonas syringae* | X | X | X | XX | X | X |
| SrG | *Stenotrophomonas sp.* | | | | | | |

**Table S3.**

**All isolates recovered from NG2 leaf wash and isolated after enrichment in M9 medium supplemented with either 4MSOB-GLS or allyl-GLS.**

| ID | family | genus | Top hit species | enriched on | Date of isolation | Used for an exp.? |
|---|---|---|---|---|---|---|
| **M1** | Microbacteriaceae | *Microbacterium* | *algeriense* | M9+allyl-GLS | 17.04.2023 | No |
| **J2** | Oxalobacteraceae | *Janthinobacterium* | *lividum* | M9+allyl-GLS | 17.04.2023 | No |
| **R3** | Yersiniaceae | not assigned (top hit: *Rahnella*) | not assigned (top hit: *contaminans*) | M9+allyl-GLS | 17.04.2023 | Yes |
| **J4** | Oxalobacteraceae | *Janthinobacterium* | *lividum* | M9+allyl-GLS | 17.04.2023 | Yes |
| **M5** | Microbacteriaceae | *Microbacterium* | *algeriense* | M9+allyl-GLS | 17.04.2023 | Yes |
| **Ps6** | Pseudomonadaceae | *Pseudomonas* | *allii* | M9+allyl-GLS | 17.04.2023 | Yes |
| **Pa7** | Erwiniaceae | *Pantoea* | *agglomerans* | M9+allyl-GLS | 17.04.2023 | Yes |
| **S8** | Xanthomondaceae | *Stenotrophomonas* | *bentonitica* | M9+allyl-GLS | 17.04.2023 | Yes |
| **Ps9** | Pseudomonadaceae | *Pseudomonas* | *lurida* | M9+allyl-GLS | 17.04.2023 | Yes |
| **MSOB1** | Pseudomonadaceae | *Pseudomonas* | *germanica* | M9+4MSOB-GLS | 07.05.2023 | Yes |
| **MSOB2** | Pseudomonadaceae | *Pseudomonas* | *germanica* | M9+4MSOB-GLS | 07.05.2023 | Yes |
| **MSOB3** | Pseudomonadaceae | *Pseudomonas* | *germanica* | M9+4MSOB-GLS | 07.05.2023 | Yes |
| **MSOB4** | Pseudomonadaceae | *Pseudomonas* | *baetica* | M9+4MSOB-GLS | 07.05.2023 | Yes |

**Table S4.**
**Primers used in this study.**

| Name | Sequence | Target / used in this study | Source |
|---|---|---|---|
| myb28_2315F | TCTTGGTCTCCGATCTATC | *myb28* gene in NG2 / Confirmation of truncation | This study |
| myb28_2316R | GCTACCTTCCATGGAAGCC | *myb28* gene in NG2 / Confirmation of truncation | This study |
| 8F | AGAGTTTGATCCTGGCTCAG | Bacterial 16S rRNA gene / Identification of 2019 isolates | (*90*) |
| 1492R | GGTTACCTTGTTACGACTT | Bacterial 16S rRNA gene / Identification of 2019 isolates | (*90*) |
| 799F | AACMGGATTAGATACCCKG | Bacterial 16S rRNA gene / Identification of 2018 isolates | (*91*) |
| 1391R | GACGGGCGGTGWGTRCA | Bacterial 16S rRNA gene / Identification of 2018 isolates | (*92*) |
| GI_At_F1 | TCCCTACACGACGCTCTTCCGATCTGTAAAGATAAATGGGTCATCTAA | *A. thaliana GI* gene / 1st PCR, amplicon sequencing | (*32*) |
| GI_At_R1 | GGAGTTCAGACGTGTGCTCTTCCGATCTTCCTTCTGAACCGGTGTATTC | *A. thaliana GI* gene / 1st PCR, amplicon sequencing | (*32*) |
| 341F-OH | TCCCTACACGACGCTCTTCCGATCTGACCTACGGGAGGCAGCAG | 16S rRNA gene / 1st PCR, amplicon sequencing | (*32*) |
| 799R-OH | GGAGTTCAGACGTGTGCTCTTCCGATCTTGCMGGGTATCTAATCCKGTT | 16S rRNA gene / 1st PCR, amplicon sequencing | (*32*) |
| Indexing_Primers_fwd | AATGATACGGCGACCACCGAGATCTACACXXXXXXXXXXXXACACTCTTTCCCTACACGACGCTCTTC | 16S rRNA gene / 2nd PCR, amplicon sequencing | Modified from (*32*) |
| Indexing_Primers_rvs | CAAGCAGAAGACGGCATACGAGATXXXXXXXXXXXXGTGACTGGAGTTCAGACGTGTGCTC | 16S rRNA gene / 2nd PCR, amplicon sequencing | Modified from (*32*) |
| At_BLc_16S_F5 | AACTTCTTTTCCAGAGAAGAAGCAAT | *A. thaliana*-specific blocking oligos / 1st PCR, amplicon sequencing | (*93*) |
| At_BLc_16S_R1 | GCTTTCGCCGTTGGTGTTCTTTCCGATCTC | *A. thaliana*-specific blocking oligos / 1st PCR, amplicon sequencing | (*93*) |

**Table S5.**

**Amplicon sequencing experiments performed as part of this study**. n.a. = not applicable.

| ID | Experiment & Figure | Leaf treat-ment | DNA extraction and purification | Plant GI gene | Primers used in 1st PCR | Blocking oligos used in 1st PCR |
|---|---|---|---|---|---|---|
| Data set 1 | Bacterial communities in minimal medium enriched with different C-sources (Fig. 5) | n.a. | SDS buffer protocol, RNAse and proteinase K clean-up, phenol/chloroform and phenol/chloroform/isoamyl alcohol clean-up, DNA precipitation | n.a. | 341F-OH 799R-OH | n.a. |
| Data set 2 | Bacterial communities in leaves of lab-grown NG2, NG*myb28*, Col-0 and *myb28/myb29* plants (Fig. 3) | Washing, or surface-sterilizing | CTAB buffer protocol, phenol/chloroform/isoamyl alcohol clean-up, DNA precipitation | Yes | 341F-OH 799R-OH | At_BLc_16S_F5 At_BLc_16S_R1 |
| Data set 3 | Bacterial communities in leaves of wild plants from the five Jena populations and co-occuring random plants, sampled in 2019 and 2020 (Fig. 4) | Washing | CTAB buffer protocol, simple DNA precipitation, magnetic bead clean-up | No | 341F-OH 799R-OH | At_BLc_16S_F5 At_BLc_16S_R1 |

**Table S6.**

**Details of the analysis of GLSs by LC-MS/MS.** GLSs were analyzed using an Agilent HPLC 1200/API3200 (AB SCIEX) instrument in negative ionisation mode. Abbreviations are: Q1, selected *m/z* of the first quadrupole; Q3, selected *m/z* of the third quadrupole; RT, retention time; DP, declustering potential (V); and CE, collision energy (V).

| **Q1** | **Q3** | **RT (min)** | **compound** | **DP** | **CE** |
|---|---|---|---|---|---|
| 436 | 95.8 | 8.6 | 4MSOB-GLS | -65 | -60 |
| 358 | 95.9 | 10.0 | Allyl-GLS | -65 | -60 |
| 388 | 95.9 | 9.0 | 2OH3Butenyl-GLS | -65 | -60 |

**Table S7.**

**Details of the analysis of amines and 4MSOB-ITC by LC-MS/MS.** 4MSOB-ITC was measured using an Agilent HPLC 1200/API3200 (AB SCIEX) instrument in positive ionisation mode. Abbreviations are: Q1, selected *m/z* of the first quadrupole; Q3, selected *m/z* of the third quadrupole; RT, retention time; DP, declustering potential (V); and CE, collision energy (V); prop-2-en-1-amine is the amine formed from allyl-GLS; 1-aminobut-3-en-2-ol is the amine formed from 2-OH-3-Butenyl-GLS.

| **Q1** | **Q3** | **RT (min)** | **compound** | **DP** | **CE** |
|---|---|---|---|---|---|
| 136 | 72 | 0.5 | 4MSOB-amine | 26 | 17 |
| 58 | 41 | 0.5 | prop-2-en-1-amine | 21 | 13 |
| 88 | 71 | 0.6 | 1-aminobut-3-en-2-ol | 21 | 13 |
| 178 | 114 | 2.6 | 4MSOB-ITC | 60 | 13 |

**Data S1.**

All strains isolated from *A. thaliana* leaves and used in this study.

**Data S2.**

Statistics for alpha diversity and beta diversity of bacterial communities in lab-grown NG2 – NG*myb28*, Col-0 – *myb28/myb29* plants (experiment for Fig. 3)
