## Supplementary material for "Beyond defense: Glucosinolate structural diversity shapes recruitment of a metabolic network of leaf-associated bacteria": Data S2

**TOTAL COMMUNITY DATA**

**Alpha Diversity (KU2t, load normalized reads, not filtered for >100 reads)**

```
# Shannon
> anova.sh = aov(richness$Shannon ~ sample_data(BacData_filt_2t)$Ecotype)
> summary(anova.sh)
                                     Df Sum Sq Mean Sq F value Pr(>F)
sample_data(BacData_filt_2t)$Ecotype  3  2.424  0.8079   1.412  0.276
Residuals                            16  9.156  0.5723
> # Chao1
> anova.ca = aov(richness$Chao1 ~ sample_data(BacData_filt_2t)$Ecotype)
> summary(anova.ca)
                                     Df Sum Sq Mean Sq F value Pr(>F)
sample_data(BacData_filt_2t)$Ecotype  3  45147   15049   1.195  0.343
Residuals                            16 201497   12594
> # Simpson
> anova.si = aov(richness$Simpson ~ sample_data(BacData_filt_2t)$Ecotype)
> summary(anova.si)
                                     Df  Sum Sq  Mean Sq F value Pr(>F)
sample_data(BacData_filt_2t)$Ecotype  3 0.02415 0.008049     0.7  0.565
Residuals                            16 0.18389 0.011493
> # ACE
> anova.ace = aov(richness$ACE ~ sample_data(BacData_filt_2t)$Ecotype)
> summary(anova.ace) # 0.0316
                                     Df Sum Sq Mean Sq F value Pr(>F)
sample_data(BacData_filt_2t)$Ecotype  3 100730   33577   3.949 0.0311 *
Residuals                            14 119034    8502
```

### Beta Diversity – Bray Curtis (KU2t, load normalized reads, filtered for >100 reads, agglomerated on genus level)

Ecotype tests all four genotypes separate

```
> adonis2(BC_Dist ~ sample_data(BacData_KU2t_gen)$Ecotype)
Permutation test for adonis under reduced model
Terms added sequentially (first to last)
Permutation: free
Number of permutations: 999

adonis2(formula = BC_Dist ~ sample_data(BacData_KU2t_gen)$Ecotype)
                                      Df SumOfSqs      R2      F Pr(>F)
sample_data(BacData_KU2t_gen)$Ecotype  3   1.0679 0.23868 1.6721  0.025 *
Residual                              16   3.4063 0.76132
Total                                 19   4.4742 1.00000
```

Glucosinolate tests WTs vs mutants

```
> adonis2(BC_Dist ~ sample_data(BacData_KU2t_gen)$Glucosinolate)
Permutation test for adonis under reduced model
Terms added sequentially (first to last)
Permutation: free
Number of permutations: 999

adonis2(formula = BC_Dist ~ sample_data(BacData_KU2t_gen)$Glucosinolate)
                                            Df SumOfSqs      R2      F Pr(>
F)
sample_data(BacData_KU2t_gen)$Glucosinolate  1   0.2981 0.06662 1.2848  0.2
18
Residual                                    18   4.1761 0.93338
Total                                       19   4.4742 1.00000
```

Wildtype tests NG+NGmyb vs. Col+myb2829

```
> adonis2(BC_Dist ~ sample_data(BacData_KU2t_gen)$Wildtype)
Permutation test for adonis under reduced model
Terms added sequentially (first to last)
Permutation: free
Number of permutations: 999

adonis2(formula = BC_Dist ~ sample_data(BacData_KU2t_gen)$Wildtype)
                                       Df SumOfSqs      R2      F Pr(>F)
sample_data(BacData_KU2t_gen)$Wildtype  1   0.5111 0.11423 2.3213  0.025 *
Residual                               18   3.9631 0.88577
Total                                  19   4.4742 1.00000
```

### Beta Diversity – Jaccard (KU2t, load normalized reads, filtered for >100 reads, agglomerated on genus level)

Ecotype tests all four genotypes separate

```
> adonis2(J_Dist ~ sample_data(BacData_KU2t_gen)$Ecotype)
Permutation test for adonis under reduced model
Terms added sequentially (first to last)
Permutation: free
Number of permutations: 999

adonis2(formula = J_Dist ~ sample_data(BacData_KU2t_gen)$Ecotype)
                                      Df SumOfSqs      R2      F Pr(>F)  
sample_data(BacData_KU2t_gen)$Ecotype  3   0.6885 0.19141 1.2625  0.082 .
Residual                              16   2.9085 0.80859                
Total                                 19   3.5970 1.00000                
```

Glucosinolate tests WTs vs mutants

```
> adonis2(J_Dist ~ sample_data(BacData_KU2t_gen)$Glucosinolate)
Permutation test for adonis under reduced model
Terms added sequentially (first to last)
Permutation: free
Number of permutations: 999

adonis2(formula = J_Dist ~ sample_data(BacData_KU2t_gen)$Glucosinolate)
                                            Df SumOfSqs      R2      F Pr(>
F)
sample_data(BacData_KU2t_gen)$Glucosinolate  1   0.3402 0.09457 1.8802  0.0
25 *
Residual                                    18   3.2568 0.90543
Total                                       19   3.5970 1.00000
```

Wildtype tests NG+NGmyb vs. Col+myb2829

```
> adonis2(J_Dist ~ sample_data(BacData_KU2t_gen)$Wildtype)
Permutation test for adonis under reduced model
Terms added sequentially (first to last)
Permutation: free
Number of permutations: 999

adonis2(formula = J_Dist ~ sample_data(BacData_KU2t_gen)$Wildtype)
                                       Df SumOfSqs     R2    F Pr(>F)
sample_data(BacData_KU2t_gen)$Wildtype  1    0.164 0.0456 0.86  0.645
Residual                               18    3.433 0.9544
Total                                  19    3.597 1.0000
```

### Beta Diversity – pairwise comparisons (KU2t, load normalized reads, filtered fo r >100 reads, agglomerated on genus level)

#### Col-0 vs. NG2

#### Bray-Curtis

```
adonis2(BC_Dist ~ sample_data(BacData_KU2t_CN)$Ecotype)
Permutation test for adonis under reduced model
Terms added sequentially (first to last)
Permutation: free
Number of permutations: 999

adonis2(formula = BC_Dist ~ sample_data(BacData_KU2t_CN)$Ecotype)
                                     Df SumOfSqs      R2      F Pr(>F)
sample_data(BacData_KU2t_CN)$Ecotype  1  0.30113 0.14296 1.3344  0.234
Residual                              8  1.80530 0.85704
Total                                 9  2.10642 1.00000
```

#### Jaccard

```
adonis2(J_Dist ~ sample_data(BacData_KU2t_CN)$Ecotype)
Permutation test for adonis under reduced model
Terms added sequentially (first to last)
Permutation: free
Number of permutations: 999

adonis2(formula = J_Dist ~ sample_data(BacData_KU2t_CN)$Ecotype)
                                     Df SumOfSqs      R2      F Pr(>F)
sample_data(BacData_KU2t_CN)$Ecotype  1   0.2566 0.15097 1.4225  0.123
Residual                              8   1.4431 0.84903
Total                                 9   1.6997 1.00000
```

#### NG2 vs. NGmyb28

#### Bray-Curtis

```
adonis2(BC_Dist ~ sample_data(BacData_KU2t_Nm)$Ecotype)
Permutation test for adonis under reduced model
Terms added sequentially (first to last)
Permutation: free
Number of permutations: 999

adonis2(formula = BC_Dist ~ sample_data(BacData_KU2t_Nm)$Ecotype)
                                     Df SumOfSqs     R2     F Pr(>F)
sample_data(BacData_KU2t_Nm)$Ecotype  1  0.53702 0.2472 2.627  0.065 .
Residual                              8  1.63541 0.7528
Total                                 9  2.17242 1.0000
```

#### Jaccard

```
adonis2(J_Dist ~ sample_data(BacData_KU2t_Nm)$Ecotype)
Permutation test for adonis under reduced model
Terms added sequentially (first to last)
Permutation: free
Number of permutations: 999

adonis2(formula = J_Dist ~ sample_data(BacData_KU2t_Nm)$Ecotype)
                                     Df SumOfSqs      R2      F Pr(>F)
sample_data(BacData_KU2t_Nm)$Ecotype  1  0.22524 0.14704 1.3791  0.042 *
Residual                              8  1.30660 0.85296
Total                                 9  1.53183 1.00000
```

Col-0 vs. myb28/29

Bray-Curtis

```
adonis2(BC_Dist ~ sample_data(BacData_KU2t_Cm)$Ecotype)
Permutation test for adonis under reduced model
Terms added sequentially (first to last)
Permutation: free
Number of permutations: 999

adonis2(formula = BC_Dist ~ sample_data(BacData_KU2t_Cm)$Ecotype)
                                     Df SumOfSqs      R2      F Pr(>F)
sample_data(BacData_KU2t_Cm)$Ecotype  1   0.2328 0.11619 1.0517  0.297
Residual                              8   1.7709 0.88381
Total                                 9   2.0037 1.00000
```

Jaccard

```
adonis2(J_Dist ~ sample_data(BacData_KU2t_Cm)$Ecotype)
Permutation test for adonis under reduced model
Terms added sequentially (first to last)
Permutation: free
Number of permutations: 999

adonis2(formula = J_Dist ~ sample_data(BacData_KU2t_Cm)$Ecotype)
                                     Df SumOfSqs      R2      F Pr(>F)
sample_data(BacData_KU2t_Cm)$Ecotype  1  0.12309 0.07136 0.6147  0.973
Residual                              8  1.60191 0.92864
Total                                 9  1.72500 1.00000
```

### ENDOPHYTIC COMMUNITY DATA

#### Alpha Diversity (KU2e, load normalized reads, not filtered for >100 reads)

```
summary(anova.sh)
                                  Df Sum Sq Mean Sq F value Pr(>F)
sample_data(BacData_filt_2e)$Ecotype  3  2.526  0.8419   1.565  0.237
Residuals                            16  8.607  0.5380
> # Chao1
> anova.ca = aov(richness$Chao1 ~ sample_data(BacData_filt_2e)$Ecotype)
> summary(anova.ca)
                                  Df Sum Sq Mean Sq F value Pr(>F)
sample_data(BacData_filt_2e)$Ecotype  3  39078   13026   1.163  0.354
Residuals                            16 179175   11198
> # Simpson
> anova.si = aov(richness$Simpson ~ sample_data(BacData_filt_2e)$Ecotype)
> summary(anova.si)
                                  Df Sum Sq Mean Sq F value Pr(>F)
sample_data(BacData_filt_2e)$Ecotype  3 0.0873 0.02911   0.838  0.493
Residuals                            16 0.5556 0.03473
> # ACE
> anova.ace = aov(richness$ACE ~ sample_data(BacData_filt_2e)$Ecotype)
> summary(anova.ace) # 0.0316
                                  Df Sum Sq Mean Sq F value Pr(>F)
sample_data(BacData_filt_2e)$Ecotype  3  12996    4332   2.672 0.0991 .
Residuals                            11  17834    1621
```

**Beta Diversity – Bray Curtis (KU2e, load normalized reads, filtered for >100 reads, agglomerated on genus level)**

Ecotype tests all four genotypes separate

```
adonis2(BC_Dist ~ sample_data(BacData_KU2e_gen)$Ecotype)
Permutation test for adonis under reduced model
Terms added sequentially (first to last)
Permutation: free
Number of permutations: 999

adonis2(formula = BC_Dist ~ sample_data(BacData_KU2e_gen)$Ecotype)
                                      Df SumOfSqs     R2      F Pr(>F)   
sample_data(BacData_KU2e_gen)$Ecotype  3   1.6242 0.2982 2.2662   0.01 **
Residual                              16   3.8225 0.7018                 
Total                                 19   5.4467 1.0000                 
```

Glucosinolate tests WTs vs mutants

```
> adonis2(formula = BC_Dist ~ sample_data(BacData_KU2e_gen)$Glucosinolate)
Permutation test for adonis under reduced model
Terms added sequentially (first to last)
Permutation: free
Number of permutations: 999

adonis2(formula = BC_Dist ~ sample_data(BacData_KU2e_gen)$Glucosinolate)
                                            Df SumOfSqs     R2      F Pr(>F
)
sample_data(BacData_KU2e_gen)$Glucosinolate  1   1.3584 0.2494 5.9809  0.00
2 **
Residual                                    18   4.0883 0.7506
Total                                       19   5.4467 1.0000
```

Wildtype tests NG+NGmyb vs. Col+myb2829

```
> adonis2(BC_Dist ~ sample_data(BacData_KU2e_gen)$Wildtype)
Permutation test for adonis under reduced model
Terms added sequentially (first to last)
Permutation: free
Number of permutations: 999

adonis2(formula = BC_Dist ~ sample_data(BacData_KU2e_gen)$Wildtype)
                                       Df SumOfSqs      R2      F Pr(>F)
sample_data(BacData_KU2e_gen)$Wildtype  1   0.1014 0.01862 0.3415   0.98
Residual                               18   5.3453 0.98138              
Total                                  19   5.4467 1.00000              
```

### Beta Diversity – Jaccard (KU2t, load normalized reads, filtered for >100 reads, agglomerated on genus level)

Ecotype tests all four genotypes separate

```
adonis2(J_Dist ~ sample_data(BacData_KU2e_gen)$Ecotype)
Permutation test for adonis under reduced model
Terms added sequentially (first to last)
Permutation: free
Number of permutations: 999

adonis2(formula = J_Dist ~ sample_data(BacData_KU2e_gen)$Ecotype)
                                       Df SumOfSqs     R2      F Pr(>F)
sample_data(BacData_KU2e_gen)$Ecotype  3   0.8686 0.2337 1.6265   0.01 **
Residual                              16   2.8481 0.7663
Total                                 19   3.7166 1.0000
```

Glucosinolate tests WTs vs mutants

```
> adonis2(J_Dist ~ sample_data(BacData_KU2e_gen)$Glucosinolate)
Permutation test for adonis under reduced model
Terms added sequentially (first to last)
Permutation: free
Number of permutations: 999

adonis2(formula = J_Dist ~ sample_data(BacData_KU2e_gen)$Glucosinolate)
                                             Df SumOfSqs     R2      F Pr(>F
)
sample_data(BacData_KU2e_gen)$Glucosinolate  1   0.5081 0.1367 2.8503  0.00
1 ***
Residual                                    18   3.2086 0.8633
Total                                       19   3.7166 1.0000
```

Wildtype tests NG+NGmyb vs. Col+myb2829

```
> adonis2(J_Dist ~ sample_data(BacData_KU2e_gen)$Wildtype)
Permutation test for adonis under reduced model
Terms added sequentially (first to last)
Permutation: free
Number of permutations: 999

adonis2(formula = J_Dist ~ sample_data(BacData_KU2e_gen)$Wildtype)
                                        Df SumOfSqs      R2      F Pr(>F)
sample_data(BacData_KU2e_gen)$Wildtype  1   0.1604 0.04317 0.8121  0.721
Residual                               18   3.5562 0.95683
Total                                  19   3.7166 1.00000
```

### Beta Diversity – pairwise comparisons (KU2t, load normalized reads, filtered for >100 reads, agglomerated on genus level)

#### NG2 vs. NGmyb28

##### Bray-Curtis

```
adonis2(BC_Dist ~ sample_data(BacData_KU2e_Nm)$Ecotype)
Permutation test for adonis under reduced model
Terms added sequentially (first to last)
Permutation: free
Number of permutations: 999

adonis2(formula = BC_Dist ~ sample_data(BacData_KU2e_Nm)$Ecotype)
                                     Df SumOfSqs      R2      F Pr(>F)
sample_data(BacData_KU2e_Nm)$Ecotype  1  0.13752 0.08852 0.7769  0.758
Residual                              8  1.41600 0.91148
Total                                 9  1.55351 1.00000
```

##### Jaccard

```
adonis2(J_Dist ~ sample_data(BacData_KU2e_Nm)$Ecotype)
Permutation test for adonis under reduced model
Terms added sequentially (first to last)
Permutation: free
Number of permutations: 999

adonis2(formula = J_Dist ~ sample_data(BacData_KU2e_Nm)$Ecotype)
                                     Df SumOfSqs      R2      F Pr(>F)
sample_data(BacData_KU2e_Nm)$Ecotype  1  0.20731 0.14221 1.3263  0.159
Residual                              8  1.25049 0.85779
Total                                 9  1.45779 1.00000
```

#### NG2 vs. Col-0

##### Bray-Curtis

```
adonis2(BC_Dist ~ sample_data(BacData_KU2e_CN)$Ecotype)
Permutation test for adonis under reduced model
Terms added sequentially (first to last)
Permutation: free
Number of permutations: 999

adonis2(formula = BC_Dist ~ sample_data(BacData_KU2e_CN)$Ecotype)
                                     Df SumOfSqs      R2      F Pr(>F)
sample_data(BacData_KU2e_CN)$Ecotype  1  0.91918 0.36943 4.6869  0.006 **
Residual                              8  1.56894 0.63057
Total                                 9  2.48813 1.00000
```

##### Jaccard

```
adonis2(J_Dist ~ sample_data(BacData_KU2e_CN)$Ecotype)
Permutation test for adonis under reduced model
Terms added sequentially (first to last)
Permutation: free
Number of permutations: 999

adonis2(formula = J_Dist ~ sample_data(BacData_KU2e_CN)$Ecotype)
                                     Df SumOfSqs      R2      F Pr(>F)
sample_data(BacData_KU2e_CN)$Ecotype  1  0.29604 0.17078 1.6476  0.074 .
Residual                              8  1.43745 0.82922
Total                                 9  1.73350 1.00000
```

**Col-0 vs. myb28/29**

**Bray-Curtis**

```
adonis2(BC_Dist ~ sample_data(BacData_KU2e_Cm)$Ecotype)
Permutation test for adonis under reduced model
Terms added sequentially (first to last)
Permutation: free
Number of permutations: 999

adonis2(formula = BC_Dist ~ sample_data(BacData_KU2e_Cm)$Ecotype)
                                     Df SumOfSqs      R2      F Pr(>F)
sample_data(BacData_KU2e_Cm)$Ecotype  1  0.12828 0.05061 0.4265  0.934
Residual                              8  2.40649 0.94939
Total                                 9  2.53478 1.00000
```

**Jaccard**

```
adonis2(J_Dist ~ sample_data(BacData_KU2e_Cm)$Ecotype)
Permutation test for adonis under reduced model
Terms added sequentially (first to last)
Permutation: free
Number of permutations: 999

adonis2(formula = J_Dist ~ sample_data(BacData_KU2e_Cm)$Ecotype)
                                     Df SumOfSqs     R2      F Pr(>F)
sample_data(BacData_KU2e_Cm)$Ecotype  1   0.1532 0.0875 0.7671  0.774
Residual                              8   1.5976 0.9125
Total                                 9   1.7508 1.0000
```
